# A mechanistic free-energy model explains and predicts sequence- and context-dependent CRISPR–Cas9 activity

**DOI:** 10.64898/2025.12.12.691775

**Authors:** Hidde S. Offerhaus, Ieva Jaskovikaitė, Stephen K. Jones, Martin Depken

## Abstract

Accurate prediction of CRISPR-based gene editing remains challenging since existing models often fail to generalize across experimental and cellular contexts. We introduce CRISPRzip, a mechanistic kinetic model that quantitatively links nucleotide sequence and environmental conditions to target interrogation. The model describes R-loop formation as movement through a sequence-dependent free-energy landscape, combining nearest-neighbor nucleic-acid energetics with protein-mediated contributions inferred from high-throughput binding and cleavage kinetics. Applying CRISPRzip to SpCas9, we predict the activity across diverse DNA targets and guide RNAs, and validate with independent singlemolecule FRET and torque spectroscopy experiments. By explicitly incorporating Cas9 concentration and DNA superhelicity, the framework predicts heterogeneous, context-dependent editing outcomes and rationalizes how physical constraints modulate cleavage dynamics. Our results provide a transferable, physics-based foundation that unifies mechanistic insight and predictive power, enabling robust characterization of CRISPR effectors and prediction of their activity across environmental contexts.

## Introduction

RNA-guided nucleases (RGNs) from CRISPR immune systems are quintessential tools for genome editing,^1^ with applications ranging from GMO crops^2^ to therapeutics.^3–5^ Their specific activity is programmed by binding a guide RNA (gRNA) that complements the intended DNA target.^6^ When the effector protein recognizes its target, it creates a double-stranded break (DSB) that can cause various edits as the cell repairs the DNA.^7–9^

Not all gRNAs produce successful edits, either because they lack activity on the intended target or because they cause undesired edits on off-target sites.^10–14^ These variable outcomes arise from a complex interplay between intrinsic factors such as the enzyme, gRNA, and target, and extrinsic factors such as delivery efficacy, cell-cycle and genomic state, and DNA repair pathways. Numerous models exist that predict the on- and off-target activity based on gRNA sequence, with the number of deep-learning models growing particularly fast.^15–19^ These models are trained in a particular cellular context, and rapidly lose predictive power at other conditions.^10,16,20^ This limited generalizability is a serious weakness given the diversity of CRISPR applications and of relevant experimental and therapeutic conditions.

Eslami-Mossalam et al. (2022)^21^ introduced a kinetic model that makes quantitative activity predictions when trained on *Streptococcus pyogenes* Cas9 (SpCas9) data. It describes Cas9 dynamics in terms of the free-energy landscapes that govern target interrogation. The underlying physical parameters can be inferred from kinetic data of Cas9 binding and cleaving a large library of DNA targets. While the model successfully predicts Cas9 activity on the basis of only mismatch positions, its insensitivity to nucleotide sequence renders it unable to capture the full variation in activity between DNA targets and RNA guides.

Here we present CRISPRzip, a general kinetic RGN framework that we apply to SpCas9, which integrates gRNA- and DNA-sequence dependence and captures variable nuclease concentrations and DNA superhelicity.

CRISPRzip incorporates the free-energy cost of the strand-replacement reaction by which Cas9 establishes an R-loop between the gRNA and the target DNA. Previous biophysical models correlated the stability of a complete R-loop with Cas9 activity.^22–25^ However, the R-loop reaction needs not equilibrate before cleavage,^26,27^ so the full kinetic pathway should be accounted for.^28^ In doing so, we capture Cas9 activity across DNA and gRNA sequences, and identify anomalous DNA targets on which Cas9:gRNA may form sequence-specific interactions beyond the R-loop hybrid.

Our predicted landscapes explain dynamics across experimental modalities, which allows us to validate CRISPRzip by comparison to Förster resonance energy transfer (FRET) assays and single-molecule torque spectroscopy measurements. The model captures the effects of Cas9:gRNA concentration and DNA superhelicity in physically meaningful parameters, and predicts and rationalizes a heterogeneous response to both these extrinsic conditions. CRISPRzip’s sequence sensitivity and ability to adapt to environments showcase its capacity to assist in gRNA design and advance CRISPR applications.

## Results

### Model definition

CRISPRzip represents the interrogation of a DNA target by the Cas9:gRNA complex in 22 partial R-loop states (Figure 1a). The complex initially binds to DNA by specifically interacting with the protospacer-adjacent motif (PAM) 5’-NGG-3’ next to a target (the ‘PAM’ state, with *b* = 0 hybrid base pairs).^29^ Next, Cas9 catalyzes the formation of an R-loop between its gRNA and the target DNA. During this process, the dsDNA separates into a displaced non-target strand (NTS)^30–32^ and a target strand (TS) that hybridizes with the gRNA. The R-loop forms in a zipper-like manner by stepwise and reversible strand replacement, starting directly upstream of the PAM. After the first dsDNA base pair is broken to form one hybrid base pair (hybrid size *b* = 1), the hybrid can either be reversed or extended. If extended, the process repeats and a longer hybrid (*b* = 2, 3, …, 20) can be formed.^32–34^ We model R-loop extension from state *b* − 1 to *b* with a single forward rate *k*_f_ (Table S2), and take R-loop recession from *b* to *b* − 1 to occur at a rate *k*_f_ exp (−Δ*U*_*b*_ / *k*_*B*_*T*), where Δ*U*_*b*_ = *U*_*b*_ − *U*_*b*− 1_ is the free-energy difference between the states. As the R-loop is completed, Cas9 undergoes a structural reconfiguration that exposes the active sites cleaving the TS and NTS.^35,36^ Experimental work shows that cleavage occurs with a rate of at least *k*_cat_ = 4.0 s^−1^.^37,38^ We simulate it as an irreversible transition from the fully opened R-loop state (*b* = 20) to the cleaved state, and adopt the above *k*_cat_ value as a model constant (Table S2). With the above parametrization, we can simulate R-loop kinetics at base pair resolution once we determine the unknown values of the free-energy landscape *U*_*b*_ = (*U*_1_, …, *U*_20_) and the R-loop extension rate *k*_f_ (Methods).

**Figure 1.**
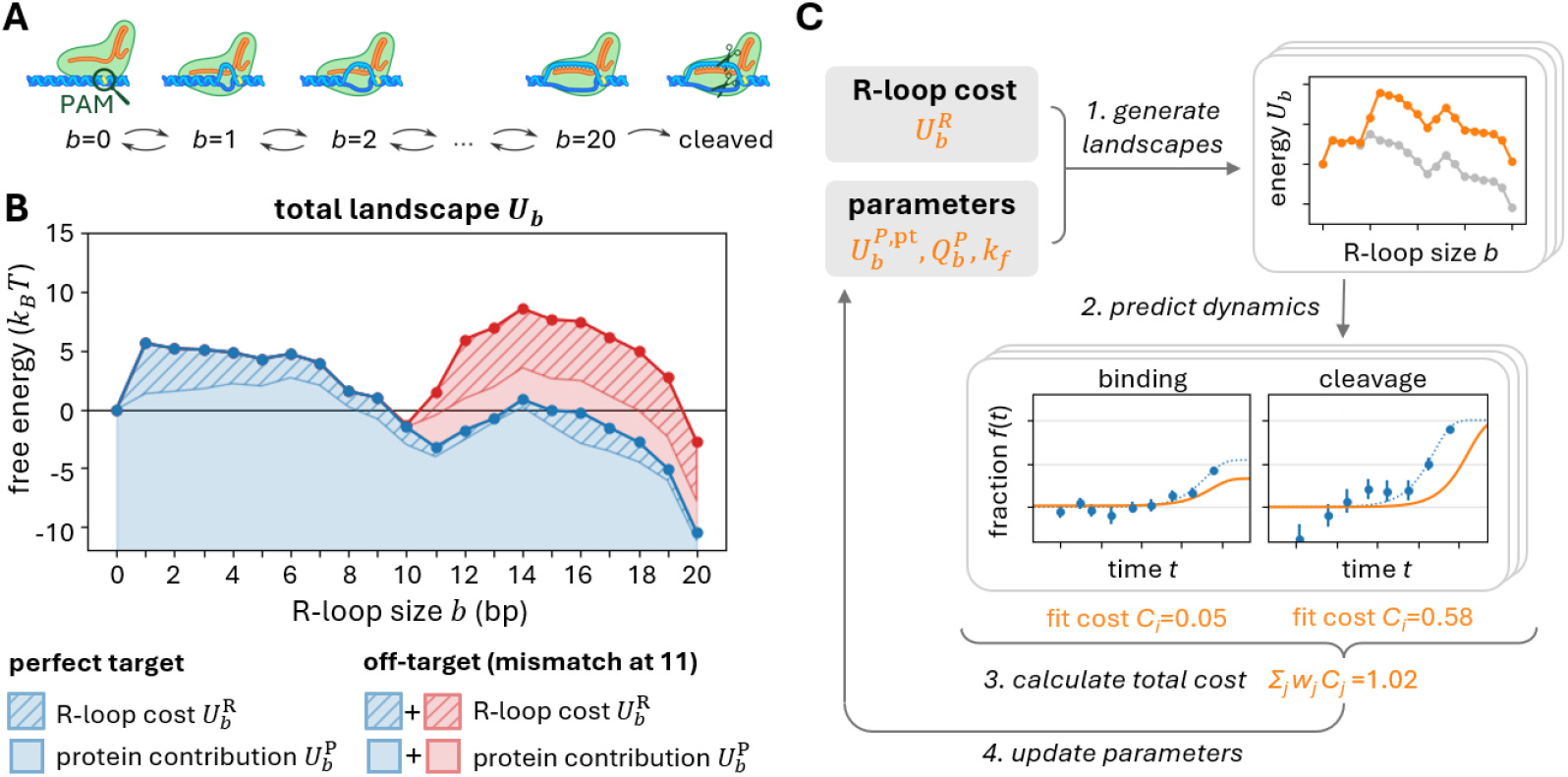
CRISPRzip model definition and training. **A** A 22-state kinetic model describes how Cas9 progressively forms an R-loop of *b* = 0, …, 20 hybrid base pairs, followed by DNA cleavage. **B** Illustration of free-energy landscape contributions by different targets. Each landscape is split up into a known R-loop cost and an unknown protein contribution, according to 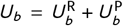, with 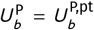 for the perfect-target landscape (blue). The landscape of the off-target (blue + red) with a mismatch at position 11 has a higher R-loop cost compared to the perfect target (red, hatched), as well as a protein penalty 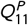 that is included in the protein contribution 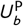 at all positions *b* ≥ 11 (red). **C** A graphic overview of the training procedure for CRISPRzip’s parameters. 1. Landscapes are created for all targets in the training set on the basis of the current set of parameters. 2. From the predicted free energy landscapes, binding and cleavage dynamics are simulated. 3. Each binding and cleavage curve is scored on its fit quality, and all scores are weighed and added up to a total cost. 4. On the basis of the total cost, the parameter set can be accepted or rejected according to simulated annealing. Steps 1–4 are repeated with newly updated parameter sets until convergence.

Our model assumes that a free-energy landscape *U*_*b*_ can be split into the sequence-dependent stability of an isolated R-loop in solution (the R-loop cost 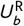), and a sequence-independent contribution due to interactions with Cas9 (the protein contribution 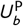, Figure 1b),

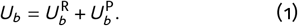

The R-loop cost 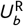 of forming an R-loop intermediate of length *b* is estimated based on nearest-neighbor parameters determined from nucleic-acid melting experiments (Figure S1, Methods).^39–43^ We follow the procedure from Alkan et al. (2018)^23^ to estimate the stability of mismatching hybrid base pairs at standard conditions. The R-loop cost at lower salt levels^44,45^ and variable temperature^39^ can be derived by uniformly scaling the standard-condition parameters by a context-dependent factor *α*.

The protein contribution 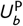 represents the total of sequence-independent interactions between Cas9 and the R-loop, including interactions with the DNA backbone and reorganizations of the protein structure that are coupled to R-loop dynamics. With a perfect (gRNA-matching) target, the protein contribution takes on the value 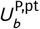. A mismatch at position *m* in the R-loop adds a protein penalty 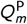 to 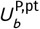 at all states *b* ≥ *m* (Figure 1b, Methods). Unlike the R-loop cost, the values of 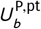 and 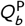 are unknown and must be inferred from experimental data (Table S2).

### Obtaining training data with high-throughput kinetic assays

We measured how DNA targets are cleaved by Cas9 and bound by catalytically dead Cas9 (dCas9). To observe gRNA sequence effects, we performed high-throughput experiments on two distinct DNA libraries derived from DNA targets λ1 and λ2 of the λ-phage genome,^46,47^ hereafter called target 1 and 2. Both libraries included the perfect target and all 1770 targets with one or two point mutations relative to the perfect target (Table S1). To measure cleavage activity, we incubated each DNA library with Cas9:gRNA ribonucleoprotein (RNP) for durations from 10 seconds to 1000 minutes (logarithmically spaced) and quantified the cleaved fraction *f*_clv_ (*t*) of each target with next-generation sequencing (NGS, Methods, Table S1). To measure binding by dCas9, we implemented a filtration assay^48^ that separated PAM-bound targets from targets bound via sufficient R-loop formation. We incubated each DNA library with RNP at the same durations as above and quantified the bound fractions *f*_bnd_ (*t*) by performing NGS on the filtrate (Methods, Table S1). Our active RNP concentration (62.5 nM) was large enough to assume saturation, ensuring that effectively each library member was Cas9-bound. Experiments were carried out at room temperature and physiological salt levels (Methods), for which we estimate an R-loop scaling factor of *α* = 0.7 ± 0.2 (Figure S2) based on DNA:RNA melting experiments at similar salt concentrations.^45^

The CRISPRzip parameters 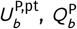 and *k*_f_ can be inferred by minimizing the total least-squared errors between our predictions and the observed bound and cleaved fractions at all measured time points (Figure 1c, Methods). The training set consists of the perfect target, all single mismatch off-targets, and 90% of the double mismatch off-targets in a library. The model performance is evaluated with a test set consisting of the excluded 10% of all double mismatch off-targets, selected by stratified sampling (Table S1).

### Establishing a single model for variable-gRNA activity prediction

Before we establish our definitive parameter values, we validate if the R-loop cost improves predictions across multiple DNA and gRNA sequences. Next, based on our training results, we construct a single parameter set that generates smooth and interpretable free-energy landscapes while maintaining performance.

Calculated R-loop costs may explain the variations among targets given a single gRNA - for instance, between two off-targets with different mismatched bases at the same position. In order to assess CRISPRzip’s capacity to capture mismatched-base effects, we set up a sequence-agnostic control model (Figure 2a). The control model’s landscapes *U*_*b*_ do not contain an R-loop cost 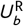, and its protein contribution 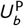 is trained to capture the average effects of mismatches based on their positions. This approach is identical to that presented in Eslami-Mossalam et al. (2022).^21^ The training results show that with no additional fit parameters, including R-loop costs reduces the total fit cost by 28% and 36% for library 1 and 2, respectively (Figure 2b). The obtained parameters (Figure S3) perform comparatively in the test sets and the training sets (Figure S4a), demonstrating their general validity.

**Figure 2.**
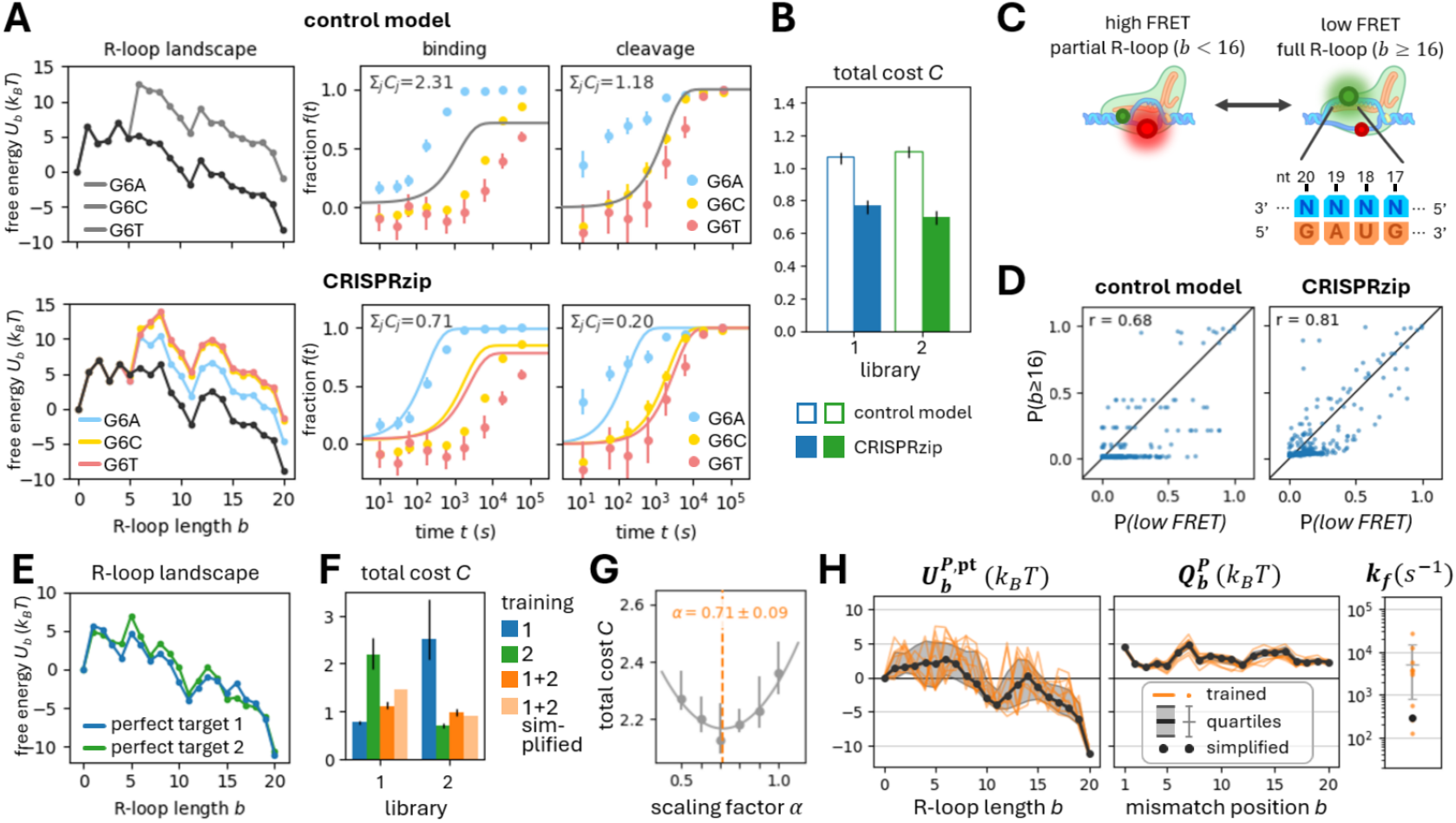
Establishing a single model for variable-gRNA activity prediction. **A** Example hybridization landscapes for the perfect target 2 (black) and three off-targets with mismatches at position 6, obtained by the reference model (top row) and with CRISPRzip (bottom row). **B** Results of training on the library 1 and 2 datasets with the control model (without R-loop cost) and CRISPRzip (with R-loop cost). Bars show average fit cost (train + test set) of the top 50% results from 200 optimization runs, error bars show 95% confidence interval. **C** Illustration of the high-throughput single-molecule FRET experiments performed by Aguirre Rivera et al.^49^ The target DNA is fluorescently labeled (Cy3 on the TS at *b* = 6 and Cy5 on the NTS at *b* = 16) such that the FRET signal drops upon R-loop completion (*b* ≥ 16). Single-molecule FRET traces are obtained for all sequence variations of the four most PAM-distal DNA bases (blue) and a single gRNA (orange) close to our gRNA 1. **D** Correlations between *P* (*b* ≥ 16), the predicted probability to attain an R-loop of size *b* ≥ 16, and *P* (*low FRET*), the measured occupancy of the low-FRET state. Dots represent the 256 DNA library members, each with a different 4-nt PAM-distal target sequence. Pearson correlation coefficient *r* is reported. **E** Joint training on library 1 and 2 predicts different R-loop landscapes for perfect target 1 (blue) and 2 (green). **F** Results of training on library 1 (blue), 2 (green), joint 1 and 2 (orange), and results of parameter simplification after joint training (light orange). Bars show average fit cost (train + test set) of the top 50% results from 200 optimization runs, error bars show 95% confidence interval. **G** Grid optimization of the R-loop scaling factor *α*. Parabolic interpolation approximates the optimum parameter value. **H** Simplification of parameters by first taking the median of 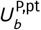 (perfect-target protein contribution) and 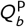 (protein penalties) from the the top-50% joint training results, and then re-optimizing the rate *k*_f_. Orange lines and dots show 10 random individual training results, shaded region and error bar show the 25%, 50%, and 75% quartiles, black dots show the simplified parameter set.

To test how far our model performance extends beyond the experimental modality used for training, we compare our models to an independent dataset of Cas9 dynamics from Aguirre Rivera et al. (2024)^49^. Here, single-molecule FRET was combined with NGS to monitor R-loop completion by Cas9 on all 256 possible sequence variations of the four most PAM-distal DNA bases (Figure 2c). The gRNA used in these experiments is nearly identical to our gRNA 1, so we use our library 1-trained parameter set to calculate R-loop formation probabilities on their DNA target library (Methods). CRISPRzip predicts the experimental results considerably better than the control model (Figure 2d), showing that model improvements persist across experimental modalities. Moreover, our Pearson correlation coefficient *r* = 0.81 approaches the correlation between repeated FRET measurements (*r* = 0.88), showing that our model explains the majority of the variation observed beyond experimental noise.

Next, we test if including the R-loop cost also allows us to predict across gRNA sequences with a single model. We train CRISPRzip on libraries 1 and 2 simultaneously, and obtain parameter sets predicting perfect-target landscapes that vary with gRNA sequence (Figures 2e and S3). The performance after joint training approaches that after isolated training (Figure 2f) and is similar for the train and test targets (Figure S4b). As a control, we apply the library 1 trained parameters to predict the library 2 data, and vice versa. This cross-prediction yields significantly worse-scoring fits (Figure 2f). Together, these observations indicate that training on multiple gRNAs increases the general applicability of the model without losing its predictive power for specific gRNAs.

Where we previously estimated the value of the R-loop scaling factor *α* on the basis of physiological-salt DNA:RNA melting experiments (Figure S2), we now co-optimize it together with the jointly-trained model parameters. We perform grid optimization at *α* ∈ [0.5, 1.0] and estimate the optimum parameter value by parabolic interpolation (Methods). The optimal R-loop scaling factor *α* = 0.71 ± 0.09 (Figure 2g) closely agrees with our previous estimate, supporting the validity of the above parameter sets.

Most joint trainings consistently reproduce local energy minima in the perfect-target protein contribution 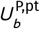 at position 0, 11 and 20 (the ‘PAM’, ‘intermediate’ and ‘open R-loop’ states), but show considerable variation elsewhere (Figure S3). As our experiments cannot resolve individual R-loop steps, the measured effective transition dynamics between the metastable states can be realized by a variety of intervening barrier regions. Among the diversely shaped barrier regions in our training results, those with high peaks are generally accompanied by large *k*_f_ values. To reveal common features across landscapes, we take the median of the model parameters 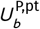 and 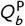 from the 50% best-scoring solutions (Figure 2h). As this median landscape is less peaked than the trained landscapes, we retrain the R-loop rate *k*_f_ (Figure 2h). The resulting simplified parameter almost completely maintains performance relative to the joint trainings (Figure 2f), only increasing the fit cost for binding double-mismatched library 1 targets (Figure S4b). As we will argue below, our model’s inability to capture the activity on these targets may arise from anomalous interactions beyond R-loop stability. All future analyses use this simplified parameter set because it makes consistent predictions while its smooth landscapes give visual insight into the effect of mismatches on R-loop dynamics.

### Free-energy landscapes explain variation in cleavage and binding dynamics

The DNA targets in our libraries show a wide variety of binding and cleavage dynamics, much of which can be understood with CRISPRzip’s predicted landscapes. The double-barrier structure of the protein contribution 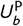 generally carries over to the total R-loop landscape 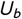. For many and diverse targets, the R-loop kinetics can be explained by simple landscape features: the height of both barriers and the stability of the intermediate and final (open R-loop) state (Figure 3a).

**Figure 3.**
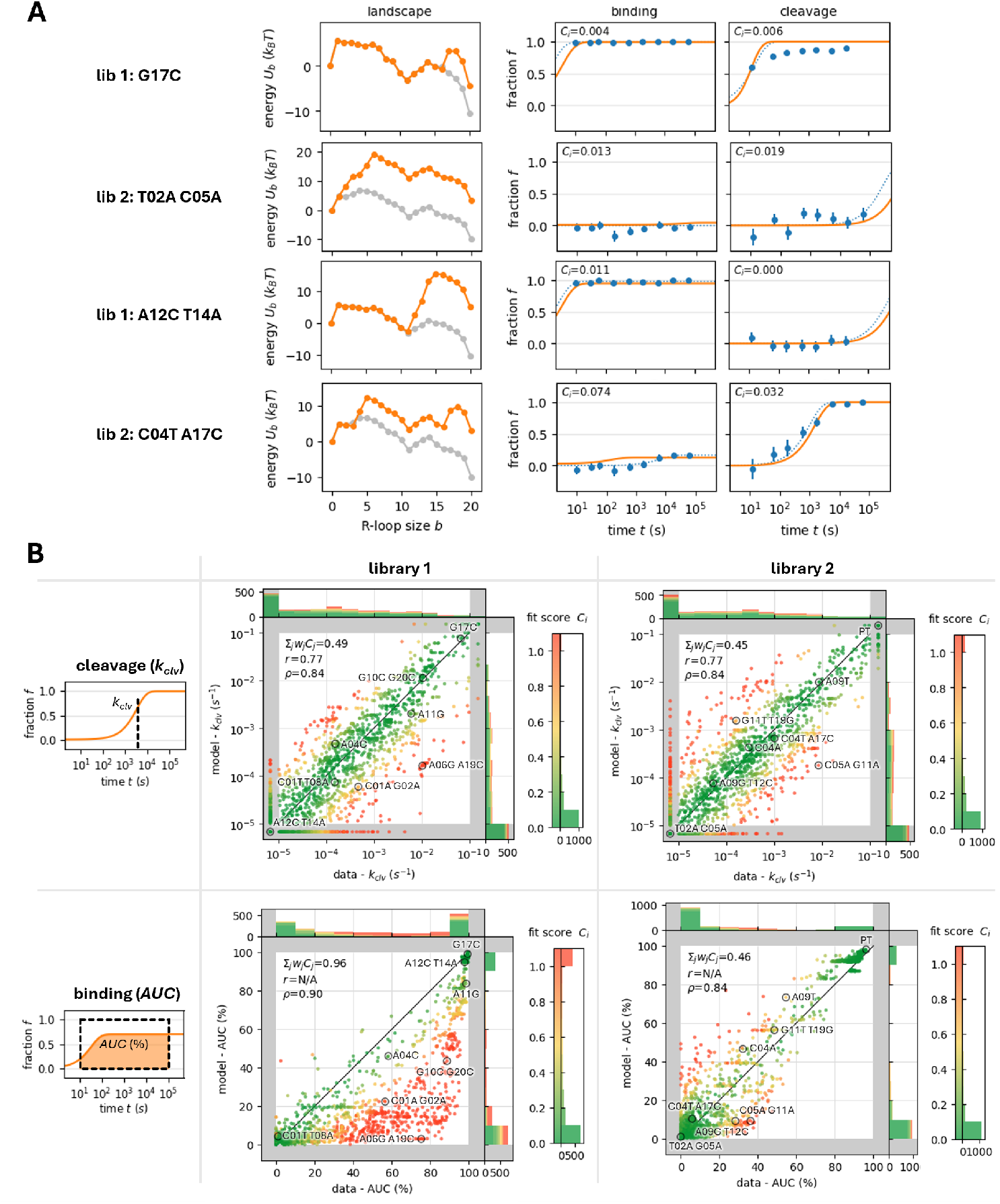
Free-energy landscapes explain variation in cleavage and binding dynamics. **A** Predicted landscapes and kinetics (orange) of several off-targets, with perfect-target landscapes (gray) for reference. Experimental data (blue) with error bars indicating standard deviation and with exponential fit curve (dotted line). Fit cost *C*_*i*_ for a target *i* is the sum of squared errors of the CRISPRzip predictions compared to that of the exponential fit curve. **B** Correlations between the kinetic data and CRISPRzip predictions. Cleavage dynamics (top row) are expressed in terms of the effective cleavage rate *k*_clv_, binding dynamics (bottom row) are summarized by the relative area under the curve (AUC, Methods). Values of *k*_clv_ outside the dynamic range [10^−5^, 10^−1^] are collapsed to the gray border region. Colors indicate fit costs *C*_*i*_ of binding or cleavage fit curves. The total weighted sum of fit costs Σ_*j*_ *w*_*j*_ *C*_*j*_ is reported for each of the four datasets. The Pearson correlation coefficient *r* is reported for the cleavage data (*k*_clv_ trimmed to the dynamic range), but not for the binding data because its near-bimodal AUC distributions violate the coefficient’s requirement of approximate normality. Spearman’s rank correlation coefficient *ρ* is reported for all data (with *k*_clv_ clipped to the dynamic range). Circles highlight the 16 selected targets for which example landscapes, binding curves and cleavage curves are shown (Figure S5).

We assess CRISPRzip’s capacity to capture the experimental variation by visualizing its activity predictions on the full target libraries (Figure 3b). The cleavage kinetics are summarized in terms of the effective cleavage rate *k*_clv_, which is fit to the experimental and simulated data (Methods). The binding kinetics are summarized in terms of the area under the curve (AUC) of fit functions to the experimental and simulated data, which reflect both the rate of binding *k*_bnd_ and the final binding level *ϕ*_bnd_ (Methods). In the Supplements, a representative selection of targets is presented in closer detail to illustrate the relation between the R-loop landscapes and the predicted binding and cleavage kinetics (Figure S5). The predictions generally correlate well to the kinetic data, reproducing cleavage and binding dynamics across a wide range of timescales. Standard correlation metrics do not accurately represent model performance, as Pearson’s *r* excludes the many *k*_clv,*i*_ values outside the dynamic range and is invalid for the bimodal AUC distributions, and Spearman’s *ρ* is dominated by boundary clusters and is insensitive to the linear relation between model and data values. Instead, the weighted total fit cost Σ_*j*_ *w*_*j*_ *C*_*j*_ provides the best indicator of quantitative agreement between predicted and experimental *k*_clv_ and AUC values.

For a group of 441 targets in library 1, making up 25% of the library, CRISPRzip consistently underestimates the binding level *ϕ*_bnd_ (Figures S6a and S6b). 92% of these targets are characterized by one of 17 specific mutations at positions 2–10 (G02C/T, C03A/T, A04T, G05A/T, A06G, G07A, T08A/C, T08G, A09C/G, G10A/C/T), followed by any mismatch at position 11– 20 (Figure S6c). Among targets with this pattern, 80% is part of the underestimated-binding group. Interestingly, we correctly reproduce the cleavage dynamics of all anomalous targets, with the exception that targets with the mutation A06G are cleaved faster than predicted. We hypothesize that these mutations stabilize the intermediate state through protein-sequence interactions not captured by CRISPRzip.

### CRISPRzip adapts to low Cas9 concentrations

Although our landscapes are trained with saturating Cas9 concentrations, we can generalize CRISPRzip to describe the dynamics at lower concentrations. We extend our 22-state model with an initial state representing an unbound DNA target. The kinetics of PAM-binding and -unbinding are described by the rates 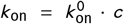 and *k*_off_ respectively, with *c* denoting RNP concentration and 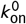 the molar PAM-binding rate. In order to determine the PAM dissociation constant 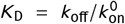, we performed cleavage assays on three library-1 targets at 7 RNP concentrations, ranging from 0.1 nM to 2 µM (Methods). We globally fit CRISPRzip predictions to the cleaved fractions and obtain a value of *K*_D_ = 29.7 ± 4.3 nM (Figure 4a). These findings are implemented in CRISPRzip with the parameters values *k* ^0^ = 3.4 · 10^−2^ nM^−1^ s^−1^ and *k*_off_ = 1.0 s^−1^ (Table S2), in line with the PAM-bound Cas9 dwell times observed in *in vitro* single-molecule FRET experiments.^34^

**Figure 4.**
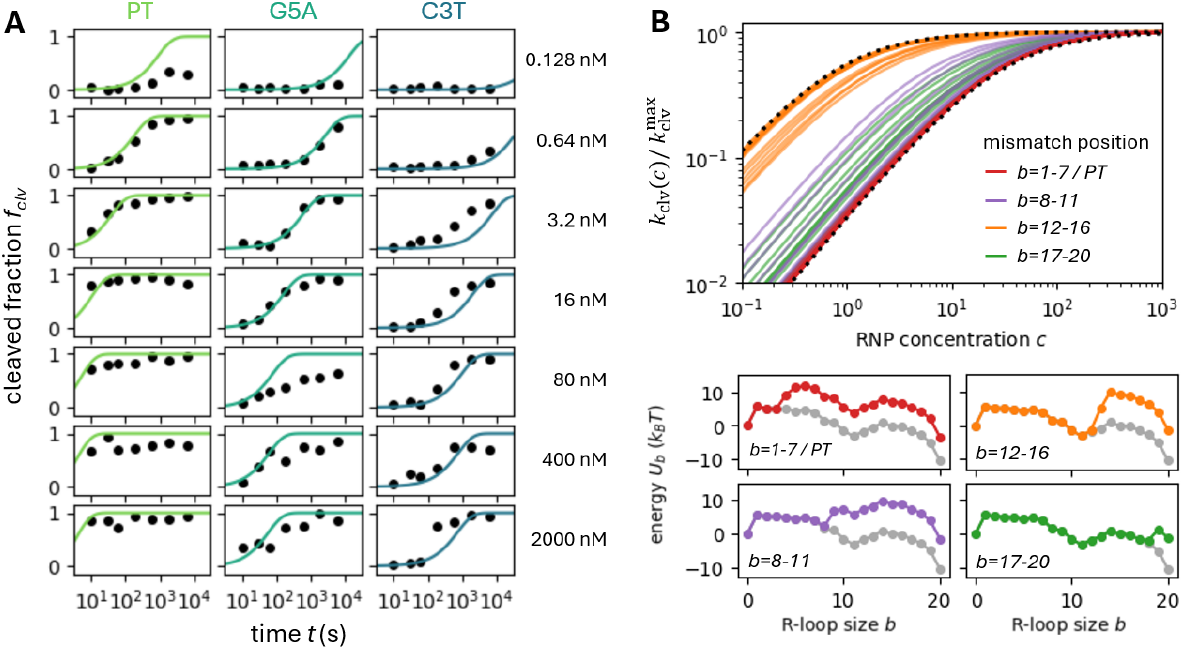
CRISPRzip adapts to low Cas9 concentrations. **A** A global fit to the cleavage of the perfect target (PT) and off-targets G5A and C3T from library 1 at various RNP concentrations *c*. Dots show data from filtration assay, lines show CRISPRzip predictions. **B** Simulated values of effective cleavage rate *k*_clv_ as a function of RNP concentration for the perfect target and all single-mismatch targets in library E. All *k*_clv_ values are shown relative to the maximum cleavage rate 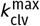 at saturating concentrations. Bottom dashed curve corresponds to Equation 2, top dashed curve shows the same relation for the effective dissociation constant 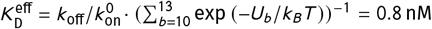· indicating the maximum intermediate state stability. Example landscapes are shown for each mismatch region.

When we predict the effective cleavage rate *k*_clv_ for all single-mismatch targets in library E, we see that Cas9 concentrations do not affect targets equally (Figure 4b). For the targets with mismatches at positions 1–7 (including PT, G5A and C3T), cleavage follows Michealis-Menten kinetics,

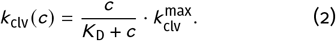

Their first landscape barriers dominate the R-loop kinetics, hence *c* modulates their cleavage rate *k*_clv_ through the level of PAM-binding. Mismatches present at positions 12–16 create a rate-limiting PAM-distal barrier, where occupancy of the intermediate state, instead of the PAM state, defines the cleavage rates. Targets with single mismatches at positions 8– 11 or 17–20 display intermediate concentration dependence as neither barrier dominates.

### CRISPRzip adapts to DNA superhelicity

DNA superhelicity alters Cas9’s capacity to cleave and bind targets.^50^ Thanks to the physical basis of CRISPRzip, we can adapt it to capture the energetic effects of DNA superhelicity on R-loop formation (Figure 5a). To simulate the effects of a torque *τ* on a DNA target, the free-energy landscape *U*_*b*_ includes the work to extend the R-loop by one base pair, which requires helix unwinding over an angular distance *θ* (Methods). The torque may affect the R-loop extension rates too, so these are updated according to the work to reach the transition state between states *b* and *b* + 1 at an angle *φ*.

**Figure 5.**
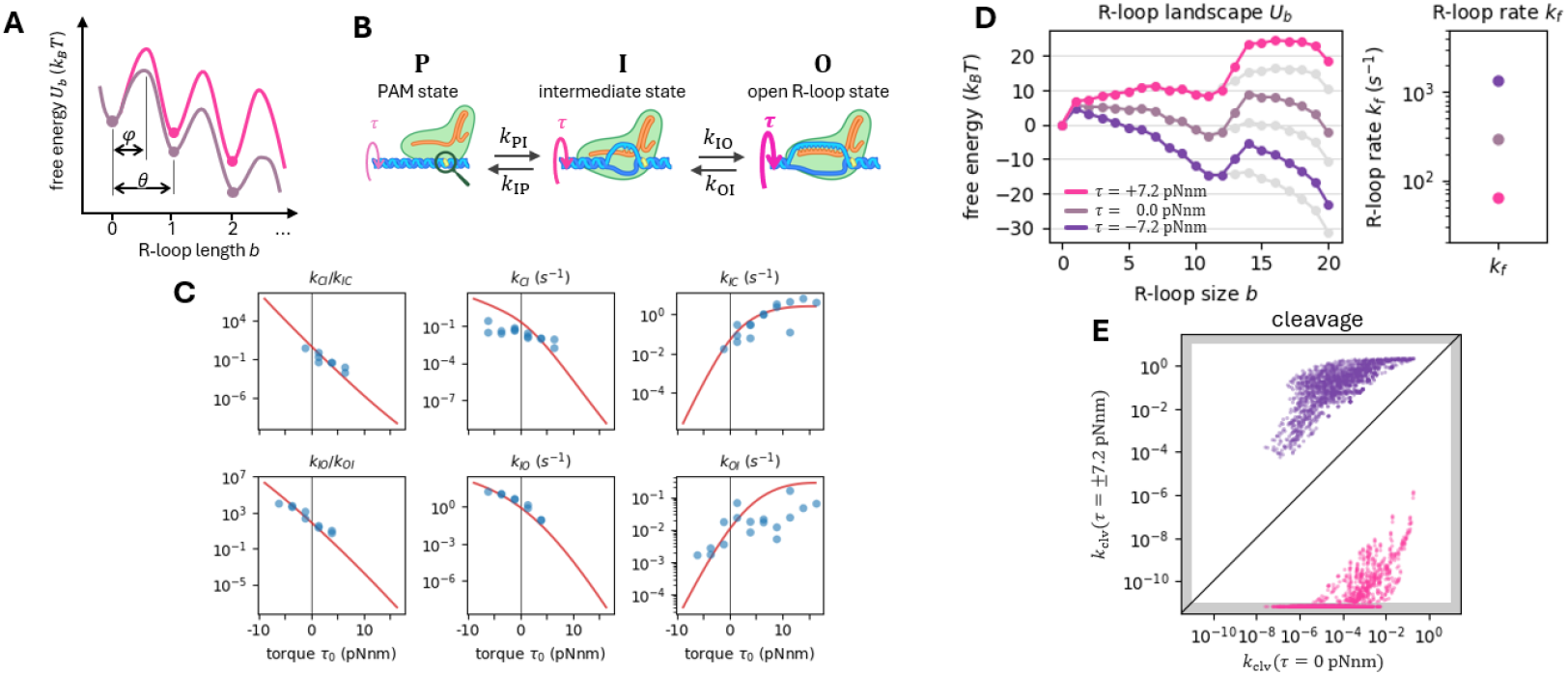
CRISPRzip adapts to DNA superhelicity. **A** Energetic effects of torque on the free-energy landscape *U*_*b*_ and its transition barriers. **B** Illustration of the observed metastable states and their effective transition rates. R-loop formation may be coupled to an increase in torque, depending on the DNA substrate. **C** Fitting the single-molecule twisting experiments of Ivanov et al. (2020)^47^. Experimental data is displayed as blue dots, the CRISPRzip fit is shown as a red line. The x-axis reports torque *τ*_0_ on unopened DNA, which increases by up to 2.8 pN nm during R-loop formation. **D** Energy landscapes *U*_*b*_ and R-loop rate *k*_f_ for perfect target 1 (light gray) and its A13T off-target at constant torque *τ* = +7.2 pN nm (pink), at zero torque (gray), and at *τ* = −7.2 pN nm (purple), with the latter corresponding to bacterial genomes at superhelical density *σ* = −0.05. **E** Predictions of cleavage activity (*k*_clv_) on all library-1 targets, comparing relaxed DNA (*τ* = 0 pN nm) to DNA under negative (*τ* = −7.2 pN nm, purple) and positive torque (*τ* = +7.2 pN nm, pink).

In Ivanov et al. (2020)^47^, magnetic tweezers experiments reveal the single-molecule kinetics of Cas9 on perfect target 1 at variable torque levels. The experiments indicate that R-loop formation requires full BDNA unwinding, yielding *θ* = 34° (Figure S7). Next, the study reports the effective rates between the PAM state ‘P’ (*b* = 0), the intermediate state ‘I’ (*b* = 11), and the open R-loop state ‘O’ (*b* = 20) (Figure 5b). When we fit our model to the measurements at variable torque (Methods), we obtain a transition state angle of *φ* = 50 ± 5°. The large value of *φ* compared to *θ* suggests that extending the R-loop requires temporary melting of the DNA also beyond the new hybrid base pair. As a consequence, the forward and backward rates between the R-loop states both slow down with positive torque and speed up with negative torque. With only one new fit parameter, CRISPRzip largely reproduces the data and captures the nonlinear dependence on torque (Figure 5c). This nonlinear effect arises from shifting the position of the highest barrier between the metastable states as torque increases (Figure 5d).

With these torsion parameters, we predict cleavage activity on the library 1 targets at an estimated bacterial genome torque of *τ* = −7.2 pN nm (Methods) and at a positive torque of *τ* = +7.2 pN nm. Our results show that torque alters cleavage rates *k*_clv_ heterogeneously (Figure 5e). For instance, negative torque accelerates the cleavage of targets with a factor 100– 30,000, depending on the relative height of the two landscape barriers.

## Discussion

CRISPRzip is a kinetic model that mechanistically captures the effects of nucleotide sequence and experimental context on RGN target recognition. By accounting for the progressive cost to form an R-loop, it quantitatively captures the variation across different DNA target and gRNA sequences when trained to SpCas9 binding and cleavage data. We validate the model across experimental modalities. CRISPRzip generates interpretable free-energy landscapes that illustrate target interrogation kinetics, and we isolate anomalous targets where Cas9 may interact specifically with the nucleic acid beyond R-loop base pairing. The environment heterogeneously affects Cas9’s cleavage dynamics, as we demonstrate for both variable Cas9 concentration and variable DNA superhelicity. Adaptation to these extrinsic factors exemplify CRISPRzip’s ability to model and predict activity in diverse applications.

Our capacity to capture sequence-specific effects relies on several factors. First, we need accurate nearest-neighbor parameters to calculate the true R-loop cost. Currently, we uniformly scale standard-condition parameters to estimate nucleic acid stability at physiological salt levels, but this simple scaling disregards the details of how base pair stabilities change with salt.[51] Optimal model performance would require that free energies of matching DNA:DNA and mismatching RNA:DNA duplexes would be determined at these conditions in nucleic acid melting experiments. Alternatively, molecular dynamics could estimate these parameters,^52,53^ and reveal how the presence of the Cas9 protein modifies the sequence-specific R-loop cost, as Cas9 bends DNA^35,54,55^, holds the gRNA-TS hybrid in A-form^31^, fixes the NTS backbone^35^ and sterically constrains the nucleic acids.

Second, to obtain the protein contributions to the free energy landscapes, we should train on a set of gRNAs and DNA targets that more broadly capture the sequence space. While our current data sets comprise many DNA sequences, it would be informative to train on more than two gRNAs, especially if they have sequence features like GC-content that differs from our gRNAs 1 and 2 (Figure S8). Previous work presented a method to quantify Cas9 binding and cleavage dynamics for many gRNAs in parallel^56^, however we could not reproduce their gRNA 1 and 2 kinetics and thus did not train with it.

Third, we need experimental verification that sequence-specific R-loop costs translate to changes in the total free energy landscape of target interrogation. Single-molecule assays probe the stabilities of semi-stable landscape states for Cas9 and other Cas nucleases,^34,47,57–59^ with recent single-molecule work revealing the full energy landscape of R-loop formation by Cascade.^60^ With these techniques, one can directly observe the influence of DNA and gRNA sequence on Cas9 kinetics, and hence provide a direct test for an energetic model of sequence effects. For instance, they could indicate if the anomalous binding patterns that we observed could arise from non-local mismatch effects.

Although the R-loop stability is a major contributor to sequence dependence, the Cas9 protein can also specifically interact with the DNA and gRNA sequence when bases form contacts with the protein structure or when the R-loop geometry is distorted.^55,61^ These interactions may sometimes dominate the target interrogation dynamics but currently lie beyond CRISPRzip’s scope. CRISPRzip may be complemented with neural networks to recognize such sequence features and correct its predictions accordingly. Other off-target features that CRISPRzip does not account for include non-canonical PAMs^25,62–64^ and off-targets with missing or added base pairs in comparison to the perfect target.^63^ These effects could be captured with a mechanistically motivated extension to the model. However, such an extension should balance parameter expansion with increased model performance and retained clarity.

With our explicit consideration of extrinsic factors and their physical effects, we contribute to a molecular understanding of the complex cellular processes that determine editing outcomes. For instance, Cas9 activity is expected to be strongly influenced by the local torsion state in a genome: transcription induces negative supercoiling,^65^ which correlates with increased Cas9 efficiency.^50^ In our upcoming experimental work, we present the target 1 and 2 libraries on supercoiled plasmids and quantify how Cas9 binds and cleaves these.^66^ CRISPRzip simulates and rationalizes these results thanks to its torque dependence. Similarly, Cas9 delivery impacts editing outcomes due to the interplay between delivery strategies and cellular transport pathways.^67^ By combining our mechanistic model with quantitative studies of cellular Cas9 delivery^68^, we expect that editing outcomes can be explained in physical terms such as the nuclear RNP concentration profile as a function of time. Lastly, Moreb and Lynch (2020)^69^ proposed that genetic context is a major determinant of activity, as many partially matching DNA sites may trap Cas9 and delay target search.^10^ When applied over a whole genome, CRISPRzip can estimate targets’ trapping capacities and predict their collective impact, again advancing insight into genome-scale phenomena from the bottom up.

In terms of activity prediction, we have previously demonstrated that universal mechanistic insight into the dynamics of Cas9 enables powerful and general off-target classification.^21^ CRISPRzip also offers Cas9 activity prediction that can be applied to any genome. It is made publicly available as a graphical user interface^70^, where users can provide gRNA and DNA sequences to inspect the predicted energy landscapes and binding and cleavage kinetics. We also provide a standalone Python package^71^, which can form a biophysical foundation for more complex machine learning models. Recent computational work^72^ demonstrated the promise of integrating kinetic models in a deep learning scheme to account for sequence-specific effects and cellular intricacies that are not feasible to capture mechanistically. Moreover, our model’s applications could extend beyond wild-type SpCas9, to other Cas nucleases^73^, engineered high-fidelity Cas9 variants^74^, relaxed-PAM variants^75^, and (d)Cas9 chimera like base and prime editors^1^, provided sufficient data is available to train the protein contribution to the free-energy landscape. We believe that CRISPRzip provides a concrete solution for reliable gRNA design, and facilitates the development of safe and effective CRISPR applications.

## Methods

### Kinetic model

CRISPRzip derives the target interrogation kinetics from the free energy landscape of R-loop formation (Figure 1a). In total, there are 23 states *b* ∈ {− 1, …, 21} that each DNA target can occupy: the unbound state (*b* = −1), PAM-bound state (*b* = 0), the partial R-loop states (*b* ∈ {1, …, 20}) and the cleaved state (*b* = 21). At saturating Cas9 concentrations, the unbound state may be omitted from the system. Given the rates between the above states, we can solve the dynamics of R-loop formation. We parametrize these rates with an approach adopted from Eslami-Mossalam et al. (2022)^21^. The rates of PAM binding (from state −1 to 0), PAM unbinding (from 0 to −1) and strand cleavage (from 20 to 21) are defined directly as *k*_on_, *k*_off_ and *k*_cat_. The PAM-binding rate 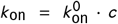 follows from the molar PAM-binding rate 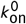 (units nM^−1^ s^−1^) and the Cas9 concentration *c* (units nM). The rates of R-loop opening and closing between states *b* ∈ {0, …, 20} are defined in terms of the free-energy landscape *U*_*b*_,

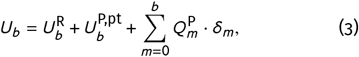

where 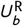 is the R-loop cost, 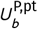 are the perfect-target protein contributions and 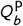 are the protein penalties. The mismatch indicator δ_*m*_ takes on the value δ_*m*_ = 0 in absence of a mismatch at position *m*, and δ_*m*_ = 1 in presence of a mismatch, such that in the latter case the penalty 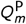 is added to all states *b* ≥ *m*. R-loop extension (from state *b* to *b* + 1) takes place at a constant rate *k*_f_. The R-loop recession rates (from *b* to *b* − 1) follow from the free energy difference Δ*U*_*b*_ = *U*_*b*_ − *U*_*b* −1_. Together, these parameters yield a Markov model described by the forward and backward rates

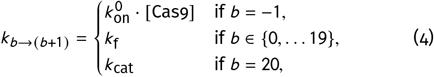

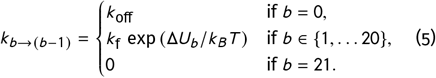

These rates form the 23 × 23 transition rate matrix **K** of the system, defined as

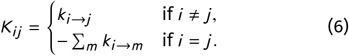

The occupancy of all states **P** (*t*) = (*P*_−1_ (*t*), …, *P*_21_ (*t*)) follows from the Master Equation *d* **P** / *dt* = **K** · **P** with the initial condition that **P** (0 = (1, 0, …, 0). The Master Equation is solved by

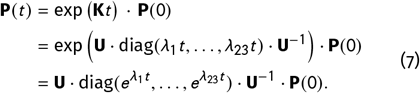

The matrix exponential is sped up by the diagonalization of the transition rate matrix **K**, writing its eigenvalues as {*λ*_1_, …, *λ*_23_} and its eigenvectors as the column vectors that make up **U**. CRISPRzip’s model definition and kinetic simulations are publicly available as a Python package.^71^

### R-loop cost

To calculate the R-loop cost, we follow the approach of Alkan et al.^23^ and sum over nearest-neighbor stability parameters for matching DNA:DNA base pairs^39^, matching^40^ and a few mismatching^41,42^ RNA:DNA base pairs. For the remaining mismatching RNA:DNA basepairs, the stabilities are estimated with an interpolation between DNA:DNA^39^ and RNA:RNA^43^ nearest-neighbor parameters. All nearest-neighbor contributions to the R-loop cost are taken at standard conditions (37 °C, 1 M NaCl), multiplied by the R-loop cost scaling factor *α*, and converted to units *k*_*B*_*T* (at room temperature *T* = 20 °C). Our calculations of intermediate R-loop costs take into account that there is a lag between DNA base stack opening and hybrid base stack formation (Figure S1); to extend an R-loop from size *b* − 1 to size *b*, the base stack between DNA bps *b* − 1 and *b* are broken up and the base stack between hybrid bps *b* − 2 and *b* − 1 is formed. Similarly, to initiate R-loop formation (*b* = 1), two DNA base stacks are broken up with no hybrid base stack in return. Our R-loop cost calculations are available in the Python package ‘crisprzip’.^71^

### Library preparation, cleavage and binding assays

The libraries 1 and 2 were designed as described in Hawkins et al. (2018)^76^ and Jones et al. (2021)^63^, and contain on-target sequences, non-target control sequences and thousands of off-target sequences with alternative PAMs or single and double mismatches, insertions, or deletions, when compared to the sgRNA. Each member has a unique barcode on each side, designed such that each cleavage product could be identified during sequencing. Each library was synthesized as pooled ssDNA oligonucleotides (GenScript), then amplified via PCR (Phusion Plus polymerase, Thermo Scientific; Table S5, primers Pr6.29/Pr6.30, Azenta Life Sciences).

For NucleaSeq cleavage experiments, described in Jaskovikaitė et al. (2025)^66^, unmodified sgRNAs (Table S4) were purchased from Synthego or GenScript, and Cas9 and dCas9 from NEB. The sgRNA, (d)Cas9 and library DNAs were diluted in 100 mM NaCl, 50 mM Tris-HCl, 10 mM MgCl_2_, 100 µg mL^−1^ recombinant albumin, pH=7.9 (r3.1 buffer, NEB). RNP complex was formed by incubating Cas9 and sgRNA at 22 °C for 15 minutes, and then library DNA was added (sgRNA:Cas9:DNA of 187.5:62.5:6.25 nM, respectively) to initiate the cleavage experiment at 22 °C. The reaction was sampled and quenched periodically (0, 0.2, 0.5, 1, 3, 10, 30, 100, 300 and 1000 min) in stop buffer (60 mM EDTA, 1 U Proteinase K, Thermo Scientific) for 30 minutes at 37 °C.

Binding affinity experiments combine elements of NucleaSeq library design and data processing with a previously described massively-parallel filter binding assay.^48,63^ The RNPs were formed and mixed with library DNA as in the cleavage experiments. The reaction was sampled at the same periodicity and stopped by passing through a 0.45 µM nitrocellulose filter (Cytiva) under vacuum.

DNA samples from cleavage and binding affinity experiments were prepared and sequenced by first appending time barcodes (indexes) with NGS prep (NEBNext, NEB), size-selecting for targets with magnetic beads (beads:DNA 0.9:1.0; AMPure XP, Beckman Coulter) and running on capillary electrophoresis (BioAnalyzer 2100, Agilent) to confirm size and purity of samples. The sample DNAs were sequenced along with 10% PhiX DNA (to ensure sequence diversity) at the EMBL GeneCore (Heidelberg, DE) on a NextSeq2000 using P2 100bp PE or P3 100bp PE chemistry.

### Determination of binding and cleavage kinetics

Read counts are determined for each library member as previously described.^63,66^ In brief, each pair of reads from the sequencing reaction are merged, and then filtered by quality, agreement between member components, and size (to separate cut and uncut DNAs). The collated read counts *c*_*j*_ (*t*) of library member *j* are expressed relative to its initial counts *c*_*j*_ (0), and are normalized to changes in library composition over time (using non-target member counts, as they are not cut by Cas9). Hence, the fraction *f*_*j*_ (*t*) of which library member *j* is bound or cleaved is calculated as

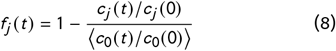

where ⟨*c*_0_ (*t*) / *c*_0_ (0)⟩ is the mean number of non-target counts relative to *t* = 0. All library members (including nontargets) with fewer than 50 initial counts are excluded from analysis. The errors in *f*_*j*_ (*t*) are estimated by assuming Poisson distributed counts *c*_*j*_ with inferred standard deviation 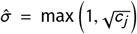 and propagating the errors from all four count terms in Equation 8. Errors of *f*_*j*_ (0) are zero by definition. When multiple library members *j* contain the same target sequence *i*, we calculate the target bound or cleaved fraction *f*_*i*_ (*t*) and its error by taking an error-weighted average of the fractions *f*_*j*_ (*t*) and their errors. Finally, we fit exponential curves to the binding and cleavage data,

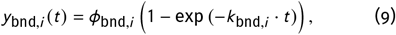

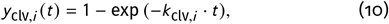

and express their fit quality in terms of residual sum of squares RSS_*i*_ = ∑_*t*_ (*y* (*t*) − *f*_*i*_ (*t*))^2^.

### CRISPRzip training

We optimize CRISPRzip’s parameters to fit the experimentally obtained binding and cleavage kinetics. The binding and cleavage data are split into a train and test set, with the test set comprising 10% of double-mismatch targets, obtained by stratified sampling of *ϕ*_bnd,*i*_ and *k*_bnd,*i*_ (binding) or *k*_clv,*i*_ (cleavage). The 41 model parameters comprise 20 perfect-target protein contribution differences 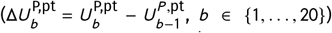 protein penalties 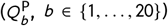 and the forward rate *k*_f_. At each optimization step, an updated set of parameters predicts bound and cleaved fractions 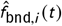 and 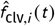 for each target *i* in the training set. The accuracy of the predictions is summarized in the cost function,

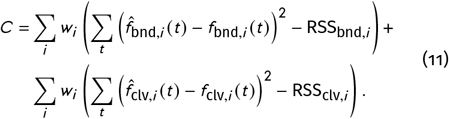

Here, the residual sum of squares between 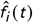 and *f*_*i*_ (*t*) are expressed relative to the RSS_*i*_ of the exponential fits *y*_*i*_ (*t*) to the data. The subtraction of these constants does not affect training but helps to interpret the contributions to cost function. The targets are weighed such that for either the set of perfect targets (a singleton), single mismatches, or double mismatches, the weights *w*_*i*_ of all member targets *i* add to 1. Given the 90/10 train-test split of double-mismatched targets, the weights are

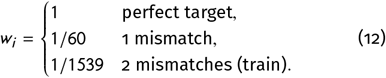

Optimization is performed with generalized simulated annealing,^77^ with initial temperature *T*_0_ = 300, visiting parameter *q*_*V*_ = 2.5, and acceptance parameter *q*_*A*_ = −5. We optimize for 1000 iterations, in each of which the parameters are changed individually in 41 update steps. We carry out 200 independent training runs on the DelftBlue high-performance computer,^78^ and select the top-50% for further analysis. For optimization of the R-loop scaling factor *α*, we carry out 25 independent training runs at fixed values *α* = 0.5, 0.6, …, 1.0 and obtain the expected optimum value by parabolic interpolation through the median fit scores.

### Binding kinetics analysis

The Area Under the Curve (AUC) is calculated for binding fit curves *y*_bnd,*i*_ (*t*) versus ln *t* in the experimental time frame *t* ∈ [*t*_1_, *t*_2_],

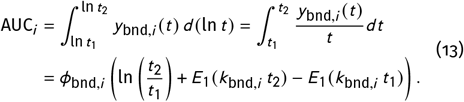

The solution is expressed in terms of the exponential integral 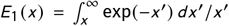. The value of AUC_*i*_ is reported relative to its maximum value, ln(*t*_2_/*t*_1_).

### Simulation of single-molecule FRET experiments

Aguirre Rivera et al. have provided us with the average FRET values for each of the 256 DNA library members in their assay.^49^ The FRET histograms in their work follow a double Gaussian distribution, *p*_*L*_𝒩(*µ*_*L*_, *σ*_*L*_) + *p*_*H*_𝒩(*µ*_*H*_, *σ*_*H*_), where the low and high FRET peaks are located at *µ*_*L*_ = 0.386 and *µ*_*H*_ = 0.690 (derived from Figure S6C), and where *p*_*L*_ and *p*_*H*_ are the weights of the low and high FRET populations, defined such that *p*_*L*_ + *p*_*H*_ = 1. Accordingly, given the average FRET signal *x*_*i*_ on a DNA target *i*, we estimate the probability to attain the low FRET state by interpolation between *µ*_*L*_ and *µ*_*H*_,

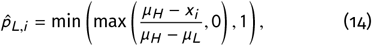

where the min and max operations guarantee probabilities in the range 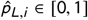.

We select the best-scoring parameter sets from our library-1 trainings of CRISPRzip and the reference model. Correcting for the two point mutations in the gRNA sequence (T instead of C, at *b* = 16 and *b* = 18), free energy landscapes are generated for each of the DNA sequences. By assuming equilibrium over all states, we obtain the state occupancies *p*_*b*_ from Boltzmann statistics,

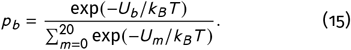

The low FRET state is chosen to correspond with R-loops of size 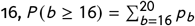, because of the position of the Cy5 label at that position. Also, the requirement that *b* ≥ 16 for low FRET signal results in a maximal correlation coefficient *r* for both the reference model and CRISPRzip predictions compared to the experimental data.

### PAM-binding experiments

Perfect target 1 (Table S5, Pr2.21, Azenta Life Sciences) and empty plasmid vector (Table S5, Pl0.63) were PCR amplified (Phusion Plus polymerase, Thermo Scientific; Table S5, primers Pr6.29/Pr6.30 and Pr6.31/Pr6.32, Azenta Life Sciences) and then column purified (GeneJET PCR purification kit, Thermo Scientific). Perfect target 1 was inserted into empty plasmid vector via HiFi assembly (NEB) and electroporated (ECM630, BTX) into homemade *E. coli* cells (TOP10 competent cells, Invitrogen). Assembled “Perfect target 1” plasmid (Table S5, Pl1.70) was purified from successful transformants (GeneJET plasmid miniprep kit, Thermo Scientific) and confirmed via sequencing (SeqVision UAB).

Two additional plasmids were made by site directed mutagenesis: one has a T instead of a C in position 3 of the target sequence (C3T), and the other has an A instead of a G in position 5 in Cas9 target sequence (G5A) (Table S5). Primers with the desired substitutions (Table S5, Pr7.77-Pr7.80) are used to PCR amplify DNA off the template plasmid (Phusion Plus Polymerase; Table S5, Pl1.70). Each reaction was exposed to DpnI to remove template (Thermo Scientific) and then electroporated into *E. coli* cells (TOP10, homemade). Several randomly picked colonies were purified (GeneJET plasmid miniprep kit, Thermo Scientific) and their sequence was confirmed (SeqVision UAB). DNA targets – perfect target, C3T, G5A – were amplified from their plasmids via PCR (Phusion Plus Polymerase; Table S5, Pr6.29, Pr6.30) and purified (GeneJET column purification kit, Thermo Scientific). Amplicon size and purity is confirmed with agarose gel electrophoresis in 1X TBE buffer (90 mM Tris-base, 90 mM boric acid, 2 mM EDTA, pH8).

The kinetics of PAM-binding and -unbinding were determined over a range of RNP concentrations (128 pM, 640 pM, 3.2 nM, 16 nM, 80 nM, 400 nM, 2 µM) This study used gRNA (Table S4) made by GenScript, diluted in 1 mM Tris, 0.1 mM EDTA, pH 8.0 upon arrival. Efficient RNP assembly was ensured by mixing Cas9 with excess sgRNA 1 (2 µM, 2.4 µM, respectively; Table S4) in r3.1 buffer, and incubating at 22 °C for 15 minutes. This RNP was also diluted to concentrations of 400 nM, 80 nM, and 16 nM. For the other concentrations, Cas9 was mixed with excess sgRNA (3.2 nM, 12.8 nM, respectively) in r3.1 buffer, and also diluted to RNP concentrations of 640 pM and 128 pM. Each RNP solution was combined with DNA at a final reaction concentration of 30 pM. The reactions were sampled and quenched periodically (0, 0.16, 0.5, 1, 3, 10, 30, 100 min) in stop buffer for 30 minutes at 37 °C. DNA cleavage was visualized on 6% Native PAGE and band intensity was quantified (GelAnalyzer 19.1, excluding background by rolling ball method, peak width tolerance = 20% of lane profile length).

### DNA superhelicity effects on Cas9 dynamics

We estimate the work done by Cas9 to unwind the DNA helix on the basis of a single molecule data from Ivanov et al. (2020)^47^. In the study, magnetic tweezers twist a linear DNA substrate containing a target that perfectly or partially matches gRNA E. A rotor bead reveals the transition rates *k*_*i j*_ between the R-loop states *i* ∈ {P, I, O}, with ‘P’ the PAM state, ‘I’ the intermediate state, and ‘O’ the open R-loop state. The torque *τ* follows from the total DNA twist *ψ* imposed by the magnetic tweezers, and the angle *ω* by which Cas9 locally unwinds DNA for R-loop formation, leading to overwinding of the tether. It is calculated as

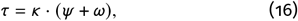

with the parameters defined such that positive and negative *ψ* and *τ* respectively reflect over- and underwound DNA, and *ω* ≥ 0 by definition. The observed torsional stiffness of *κ* = 1.48 pN nm rot^−1^ (Figure S7) is expected for a 5 kb tether with a twist persistence length *C* = 100 nm for linear DNA under a tension of 5 pN.^79,80^ R-loop completion corresponds to full B-DNA unwinding, *ω* = 681° ≈ 20 × 34° (Figure S7).

In CRISPRzip, we assume constant DNA helix unwinding with angular distances *φ* to the transition states and *θ* between the R-loop states (Figure 5a). For DNA under a torque *τ*_0_ = *κ* · *ψ* in its unopened state, the free-energy landscape *U*_*b*_ (*τ*_0_) can be obtained from the landscape *U*_*b*_ (0) on unconstrained DNA by including the angular work to extend the R-loop,

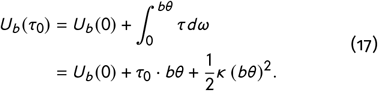

Similarly, the R-loop extension rates *k*_*b* → *b* + 1_ change according to the work to reach the next transition state,

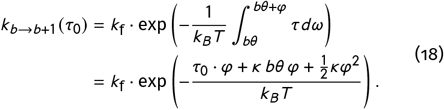

If the DNA substrate has significant torsional stiffness *κ*, the R-loop extension rates are no longer constant across all states *b*.

We fix the value *θ* = 34° on the basis of the observed full B-DNA unwinding. The transition angle *φ* is estimated by fitting transition rates 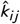 on perfect target 1 to experimental data (measured with twist cycling of 3 rpm and 15 rpm). For the PAM-intermediate transitions (*k*_PI_ and *k*_IP_), kinetic data is added from the off-target with mismatches from position *b* = 13 on. To calculate 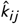 in CRISPRzip, we 1) assume a position *b* of state ‘I’; 2) calculate the arrival time distributions between states ‘P’, ‘I’ and ‘O’; and 3) fit exponential functions to these distributions with fit rates 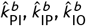 and 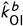. This process is carried out for assumed intermediate states *b* ∈ {9, …, 13}, after which we determine the final effective rates according to Boltzmann statistics of the intermediate states,

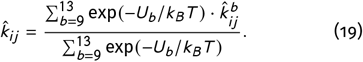

We perform a log-least-squares fit of the transition angle *φ* to the experimental rates *k*_PI_, *k*_IP_, *k*_IO_ and *k*_OI_ as well as their ratios *k*_PI_ *k*_IP_ and *k*_IO_ *k*_OI_, the latter of which are added to assign extra weight to the relative state stabilities in the optimization.

In a genome, the torsion on DNA follows from the super-helical density *σ*, which quantifies how much the helix is overwound (*σ* > 0) or underwound (*σ* < 0) compared to its relaxed state. For an *E. coli* genome with *σ* = −0.05, the DNA helix makes 5% fewer turns than it would at its natural helix repeat of *h*_0_ = 10.5 bp = 3.6 nm. The torque on a genome is given by *τ* = *C*_eff_ 2*π σ k*_*B*_*T* / *h*_0_, where *C*_eff_ = 20 nm is the effective twist persistence length of plectonemic, tensionless DNA.^79,80^ These values predict a torque of *τ* = −7.2 pN nm on supercoiled bacterial genomes at *σ* = −0.05. Due to the large sizes of genomes (5 Mb for *E. coli*), their torsional stiffness is negligibly small (*κ* ≈ 10^−4^ *k*_*B*_*T* rot^−1^) and a constant torque *τ* = *τ*_0_ can be assumed.

## Author contributions

H.S.O. and M.D. conceived and designed the study. H.S.O. developed the methodology, implemented the software, trained the model, validated the model, performed the analysis, and generated the visualizations. I.J. and S.K.J. developed the experimental protocols. I.J. performed the experiments. H.S.O. and S.K.J. curated the data. H.S.O. and I.J. wrote the original draft manuscript. H.S.O., I.J., S.K.J., and M.D. reviewed and edited the manuscript. M.D. supervised the project.

## Acknowledgements

We thank Sebastian Deindl, Anton Sabantsev and Klaus Brackmann for their involvement in the project, for their experiments on salt conditions, and for supplying their MUSCLE data^49^ and discussing how to model these. We thank Ralf Seidel, Julene Madariaga-Marcos and Fabian Welzel for our many valuable discussions about target interrogation dynamics and the implementation of nearest-neighbor models. We thank Ard Louis and Eryk Ratajczyk for our discussion on R-loop energetics and the influence of Cas9 on R-loop stability. We thank the EMBL sequencing facility (GeneCore, Heidelberg, DE) for sequencing assistance. Finally, we thank Raúl Ortiz-Merino, Elviss Dvinskis, and the TU Delft Digital Competence Centre for providing assistance in the design and development of the CRISPRzip Python package and the CRISPRzip GUI. H.S.O. acknowledges financial support from the Biononscience Department at TU Delft. This work was partially funded by the European Research Council (Project 101078247—PROTEGE; S.K.J.), the Ministry of Education, Science and Sports of the Republic of Lithuania (Program “University Excellence Initiatives” 12-001-01-01-01, project No. S-A-UEI-23-10; S.K.J.), and a European Molecular Biology Organization Installation Grant (Project IG 5728-2024; S.K.J.).

## Data and software availability

The CRISPRzip source code (v1.2.2)^71^ is available on GitHub (https://github.com/hiddeoff/crisprzip) and on PyPi (https://pypi.org/project/crisprzip/). The CRISPRzip GUI (v1.0.0)^70^ is available on GitHub (https://github.com/hiddeoff/crisprzip-tool). The NucleaSeq pipeline for processing cleavage assay read counts^63^ is available on Github (https://github.com/JonesLabEU/nucleaseq). We provide the target sequences, experimental cleavage and binding kinetics, and CRISPRzip-simulated cleavage and binding kinetics for libraries 1 and 2 as supplementary data. Raw sequencing data will be made available through the European Nucleotide Archive database.

## Supplementary Figures

**Figure S1.**
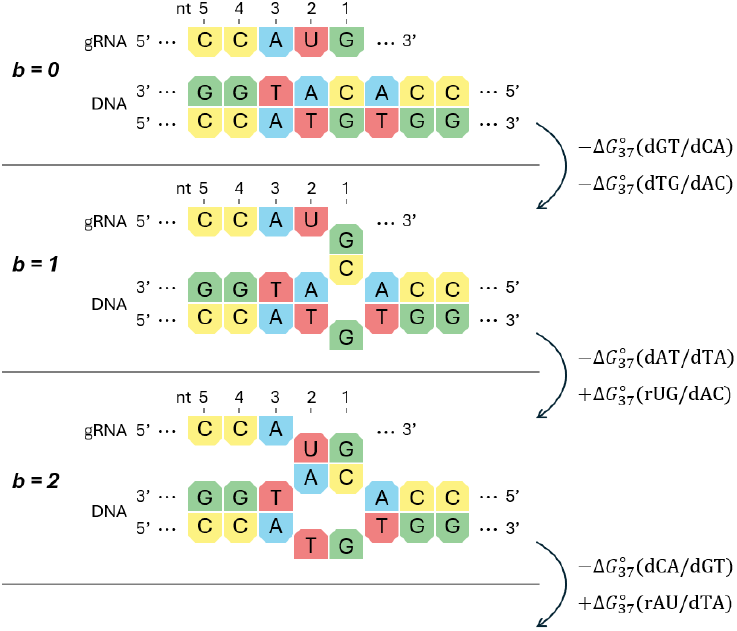
Calculation of the R-loop cost. At each R-loop extension step, a DNA:DNA base pair is broken up and an RNA:DNA base pair is established. The nearest neighbor parameters describe the stability of a stack of two neighboring base pairs, e.g. the base stack 5’-GT-3’ / 3’-CA-5’ (or equivalently, 5’-AC-3’ / 3’-TG-5’) at position *b* = 1 has a free energy 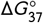 (dGT/dCA). Alkan et al. (2018)^23^ prescribes a collection of nearest-neighbor parameters and a methodology to add them, accounting for the presence of mismatches between the DNA and gRNA sequence. The total R-loop cost under standard conditions is calculated as a progressive sum over all nearest-neighbor parameters.

**Figure S2.**
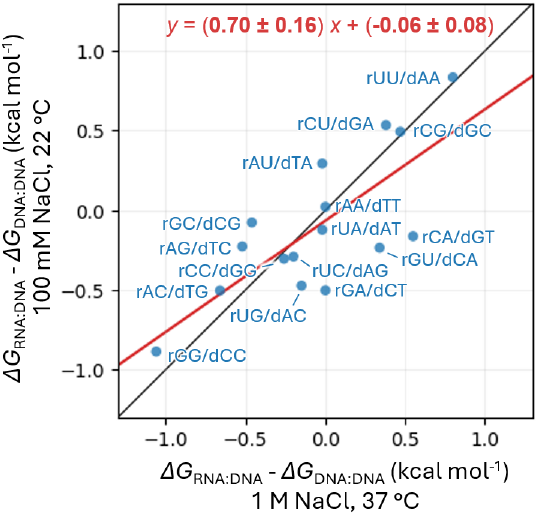
Influence of temperature and salt on the base stack exchange cost. The base exchange cost Δ*G*_RNA:DNA_ − Δ*G*_DNA:DNA_ represents the work required to replace a dsDNA base stack by a hybrid RNA:DNA base stack. No mismatching base pairs are included. The notation rGU/dCA indicates replacing 5’-GT-’3 / 3’-CA-5’ dsDNA base pairs with 5’-GU-3’ / 3’-CA-5’ hybrid base pairs. Parameters at standard conditions (1 MNaCl, 37 °C) are from SantaLucia and Hicks (2004)^39^ (dsDNA) and Sugimoto et al. (1995)^40^ (RNA:DNA). Parameters at physiological salt (100 mM NaCl) and room temperature (22 °C) are from SantaLucia and Hicks (2004)^39^ and Banerjee et al. (2020)^45^.

**Figure S3.**
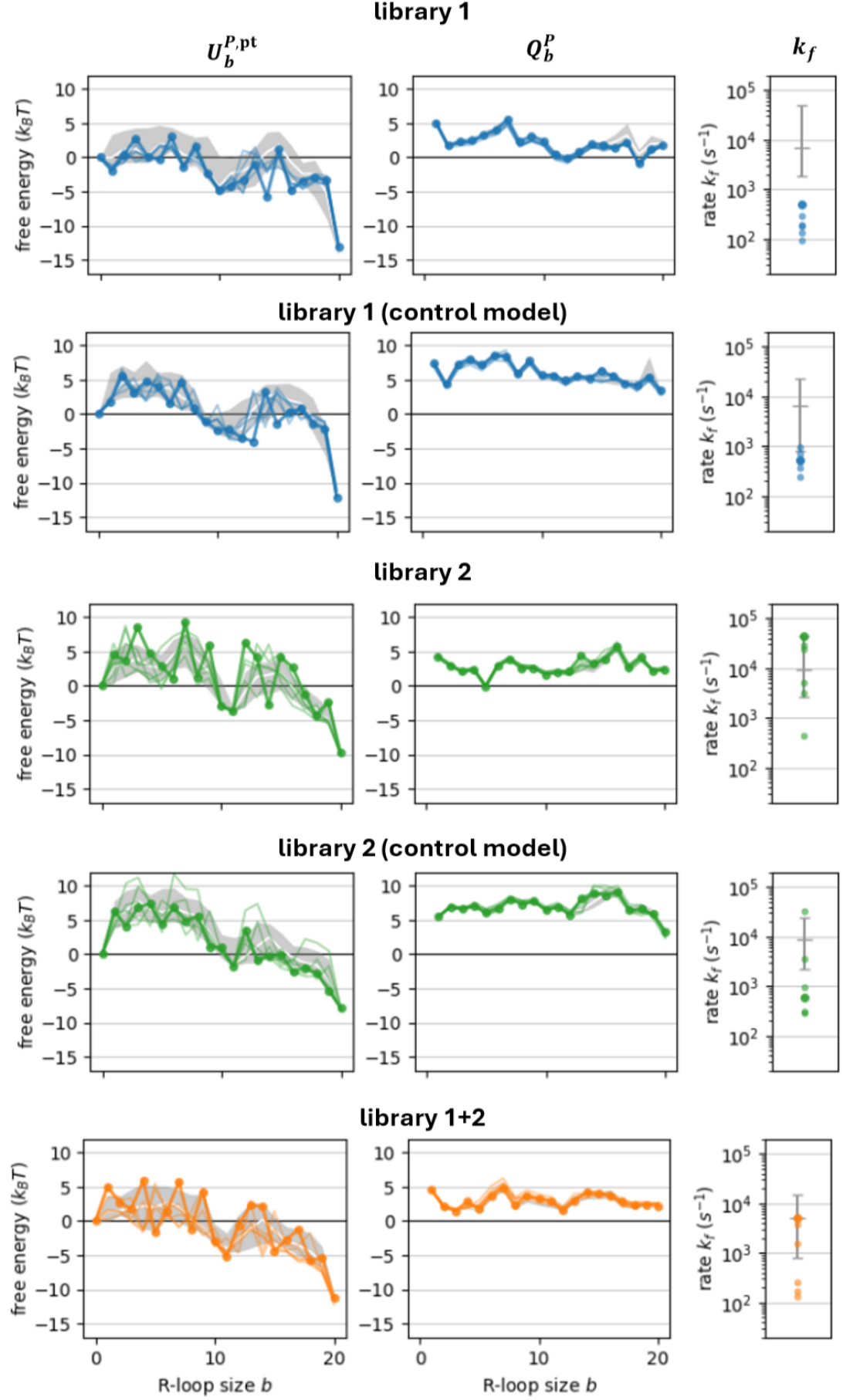
Obtained parameter sets. Example parameter sets as obtained from different training settings. For each, 200 optimization runs are performed at fixed *α* = 0.7. The parameter sets consist of 20 perfect-target protein contributions 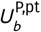, 20 protein penalties 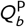 and the R-loop rate *k*_f_. Solid, dotted line shows best-scoring parameter set, light lines show the 5 next-best scoring parameter sets (shown as large and small dots in the right panels). White line indicates median of the top 50% solutions, light gray region shows their 50% confidence intervals (shown as error bars in the right panels).

**Figure S4.**
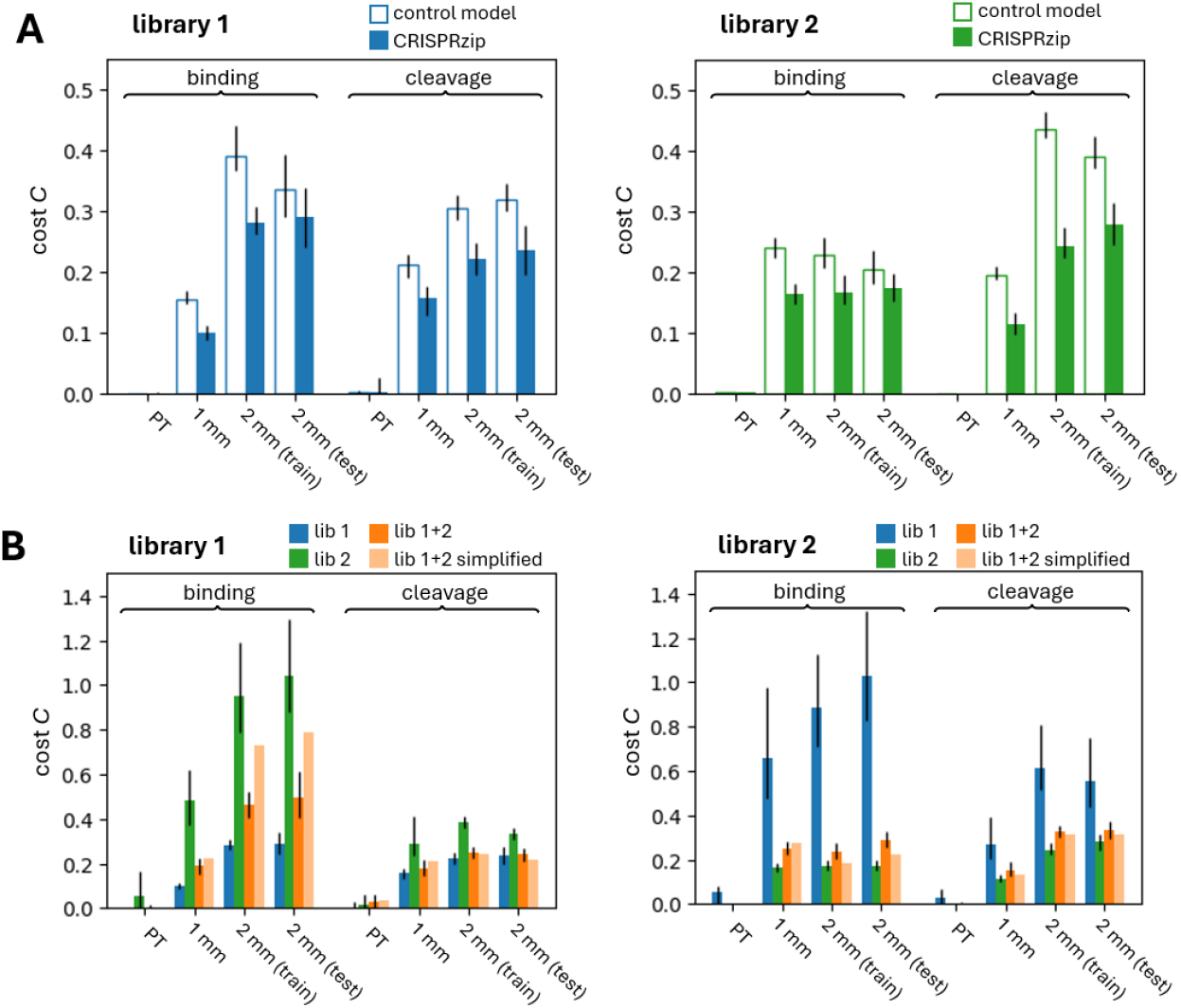
Fit scores on data subsets. **A** Partial fit costs *C* on the perfect target (PT), single mismatch offtargets (1 mm) and double mismatch off-targets (2 mm), split according to the train and test group (90%/10% data split). Bars show average fit cost of the top 50% results from 200 optimization runs, error bars show 95% confidence interval. A sum of all partial fit costs *C* weighted according to their group size (Equation 12) results in the total fit score (Figure 2a). **B** Total and partial fit costs of the top-50% parameter sets from training on library 1 and 2 in isolation (blue, green) and simultaneously (orange), together with the performance of the simplified parameter set (Figure 2a).

**Figure S5.**
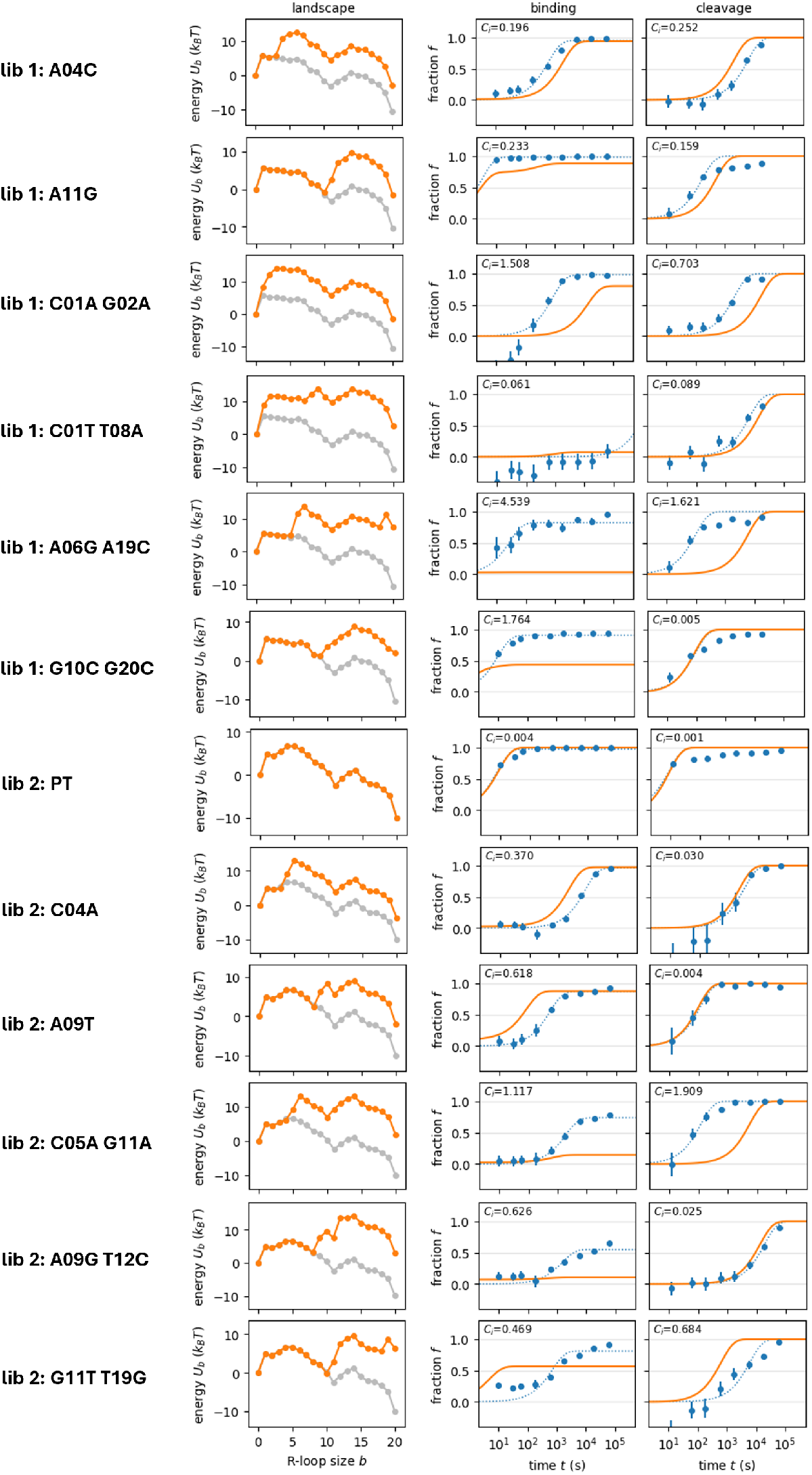
CRISPRzip predictions. Predicted landscapes and kinetics (orange) for several targets in libraries 1 and 2, made with the simplified parameter set. Perfect target landscapes (light gray) are shown for reference. Experimental data shown (blue) with error bars indicating standard deviation and with exponential fit curve (dotted line). Curve fit scores *C*_*i*_ reported next to binding and cleavage predictions.

**Figure S6.**
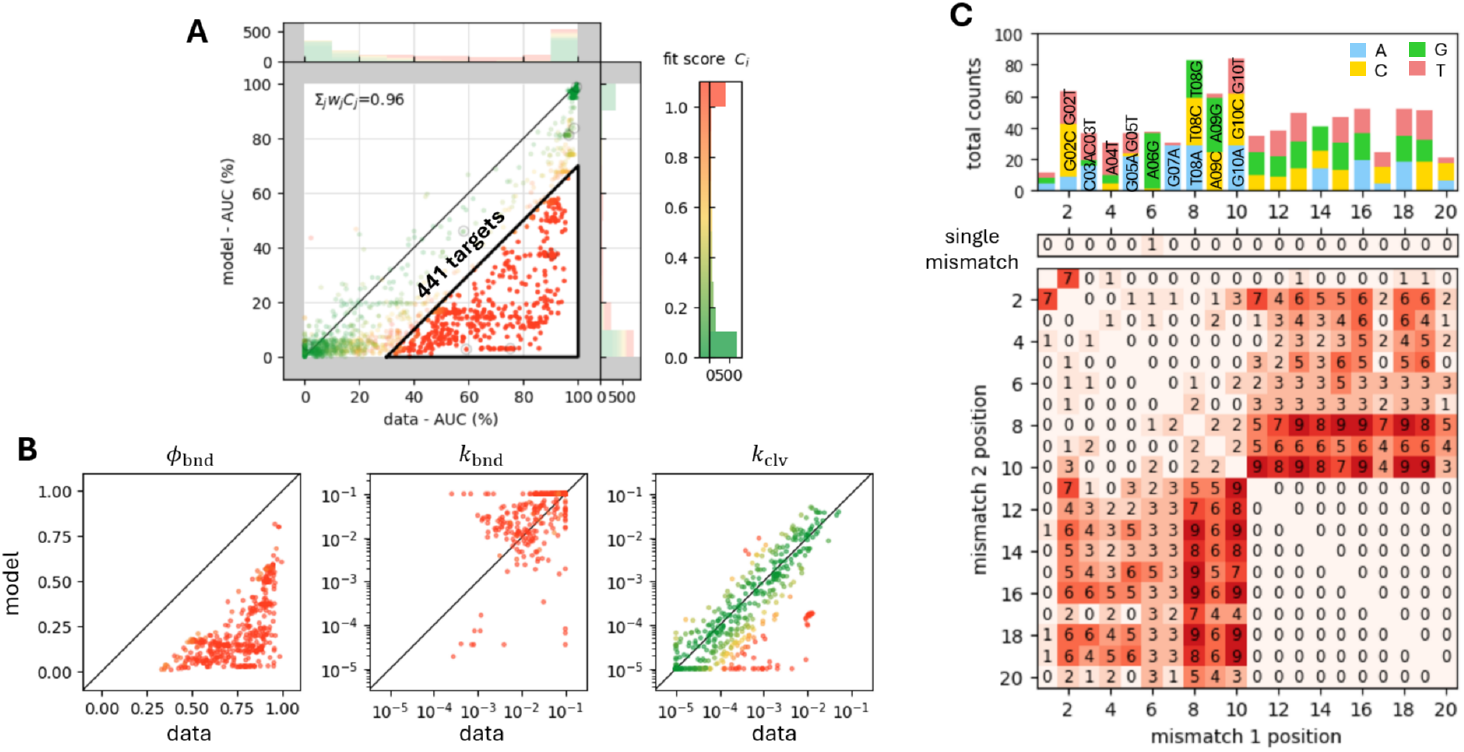
Anomalously binding library 1-targets. **A** Selection of 441 targets in library 1 that display anomalously stable binding. **B** Correlating the parameters of exponential fit curves to the experimental and simulated data: the maximum bound fractions *ϕ*_bnd_, binding rates *k*_bnd_ and cleavage rates *k*_clv_. **C** Inspection of the sequences of the 441 targets. Top panel: total occurrence of specific mismatches, by base type. Center and bottom panel: occurrence of mismatch positions, in isolation (center) or in pairs (bottom). The selection includes at most 3 targets with a single mismatch at a particular position (center) and at most 9 targets with a mismatch pair at a particular position (bottom). See Figure S5 for targets A06G A19C and G10C G20C.

**Figure S7.**
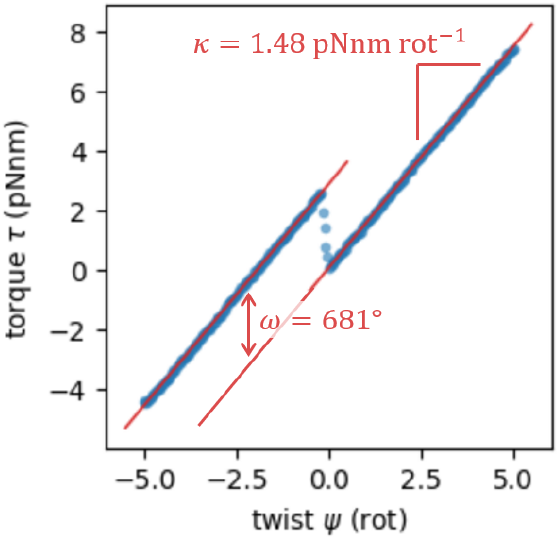
Torsional stiffness in single-molecule twisting experiments. The torsional stiffness *κ* = *τ* / *ψ* of the full DNA tether is determined with a linear fit to the torque-twist curve.

**Figure S8.**
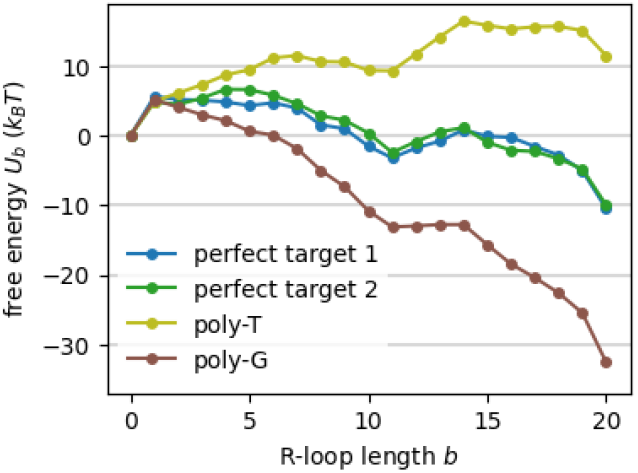
Lower and upper bounds to R-loop stability. Free energy landscapes on the most stable (poly-G target) and least stable (poly-T target / poly-U gRNA) R-loops according to nearest-neighbor parameters. Target-1 and -2 landscapes are shown for reference.

## Supplementary Data

**Table S1.** Target sequences, cleavage and binding data, and CRISPRzip predictions. See Supplemental Files.

| parameter(s) | name | value | source |
| --- | --- | --- | --- |
| $U_1^{P,pt}, \dots, U_{20}^{P,pt}$ | perfect-target protein contribution | free | |
| $Q_1^P, \dots, Q_{20}^P$ | protein penalties | free | |
| $U_1^R, \dots, U_{20}^R$ | R-loop cost | sequence-dependent | Alkan et al. (2018)<sup>23</sup> |
| $\alpha$ | R-loop scaling coefficient | 0.7 | This work |
| $k_f$ | R-loop extension rate | free | |
| $k_{cat}$ | catalysis rate | $4.0 \text{ s}^{-1}$ | Gong et al. (2018)<sup>37</sup> |
| $k_{on}^0$ | PAM-binding rate | $3.4 \cdot 10^{-2} \text{ nM}^{-1} \text{ s}^{-1}$ | This work |
| $k_{off}$ | PAM-unbinding rate | $1.0 \text{ s}^{-1}$ | This work |
| $\theta$ | R-loop step angle | $34^\circ$ | Ivanov et al. (2020)<sup>47</sup> |
| $\varphi$ | transition angle | $50^\circ$ | This work |

**Table S2.** Free and fixed parameters in CRISPRzip.

| parameter | value ($k_B T$) | parameter | value ($k_B T$) | parameter | value ($\text{s}^{-1}$) |
| --- | --- | --- | --- | --- | --- |
| $U_1^{P,pt}$ | 1.344 | $Q_1^P$ | 4.445 | $k_f$ | 292.6 |
| $U_2^{P,pt}$ | 1.546 | $Q_2^P$ | 2.097 | | |
| $U_3^{P,pt}$ | 1.766 | $Q_3^P$ | 1.548 | | |
| $U_4^{P,pt}$ | 2.194 | $Q_4^P$ | 2.048 | | |
| $U_5^{P,pt}$ | 1.990 | $Q_5^P$ | 1.750 | | |
| $U_6^{P,pt}$ | 2.697 | $Q_6^P$ | 3.657 | | |
| $U_7^{P,pt}$ | 2.065 | $Q_7^P$ | 4.839 | | |
| $U_8^{P,pt}$ | 0.225 | $Q_8^P$ | 2.566 | | |
| $U_9^{P,pt}$ | -0.818 | $Q_9^P$ | 3.237 | | |
| $U_{10}^{P,pt}$ | -2.991 | $Q_{10}^P$ | 2.816 | | |
| $U_{11}^{P,pt}$ | -4.022 | $Q_{11}^P$ | 2.703 | | |
| $U_{12}^{P,pt}$ | -2.582 | $Q_{12}^P$ | 1.851 | | |
| $U_{13}^{P,pt}$ | -1.101 | $Q_{13}^P$ | 3.164 | | |
| $U_{14}^{P,pt}$ | 0.193 | $Q_{14}^P$ | 3.691 | | |
| $U_{15}^{P,pt}$ | -1.401 | $Q_{15}^P$ | 3.625 | | |
| $U_{16}^{P,pt}$ | -2.889 | $Q_{16}^P$ | 3.910 | | |
| $U_{17}^{P,pt}$ | -3.552 | $Q_{17}^P$ | 2.382 | | |
| $U_{18}^{P,pt}$ | -4.464 | $Q_{18}^P$ | 2.282 | | |
| $U_{19}^{P,pt}$ | -6.045 | $Q_{19}^P$ | 2.444 | | |
| $U_{20}^{P,pt}$ | -11.154 | $Q_{20}^P$ | 2.233 | | |

**Table S3.**
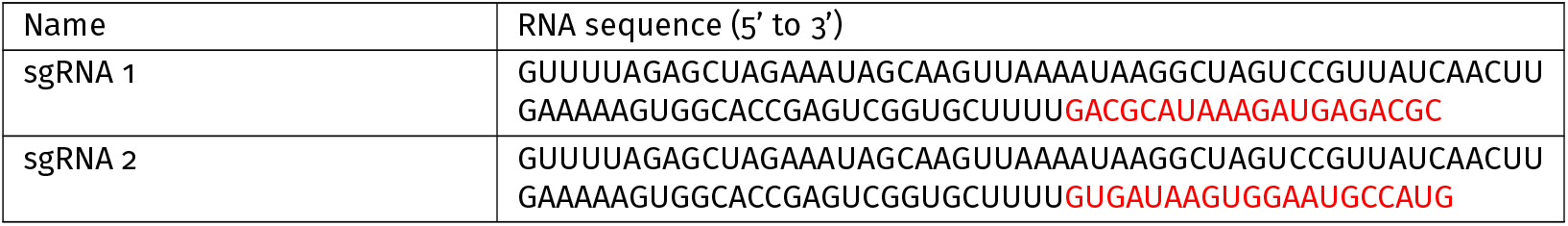
Simplified parameter set.

**Table S4.**
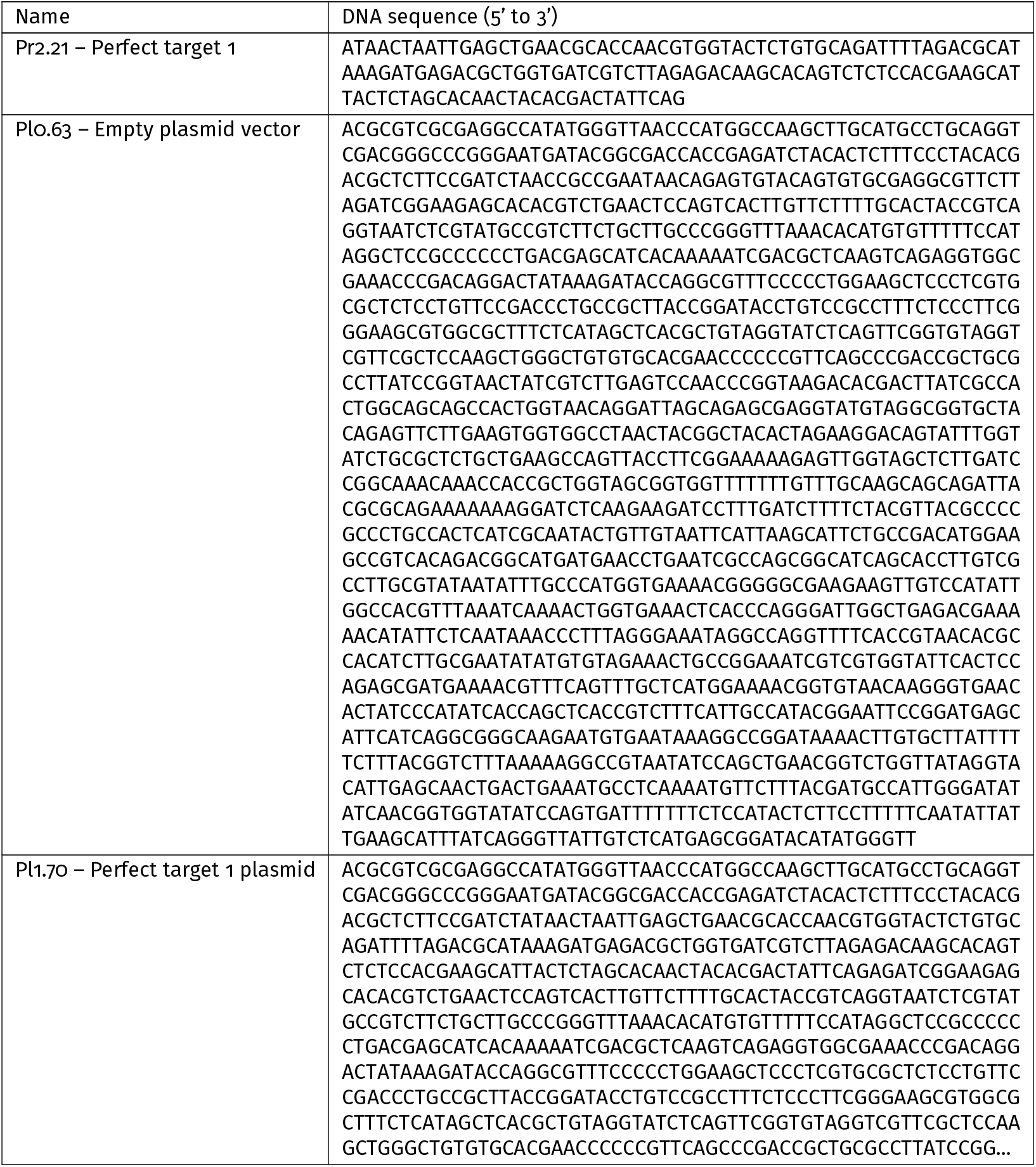
gRNA used in the study. Target-matching sequences are highlighted in red.

**Table S5.**
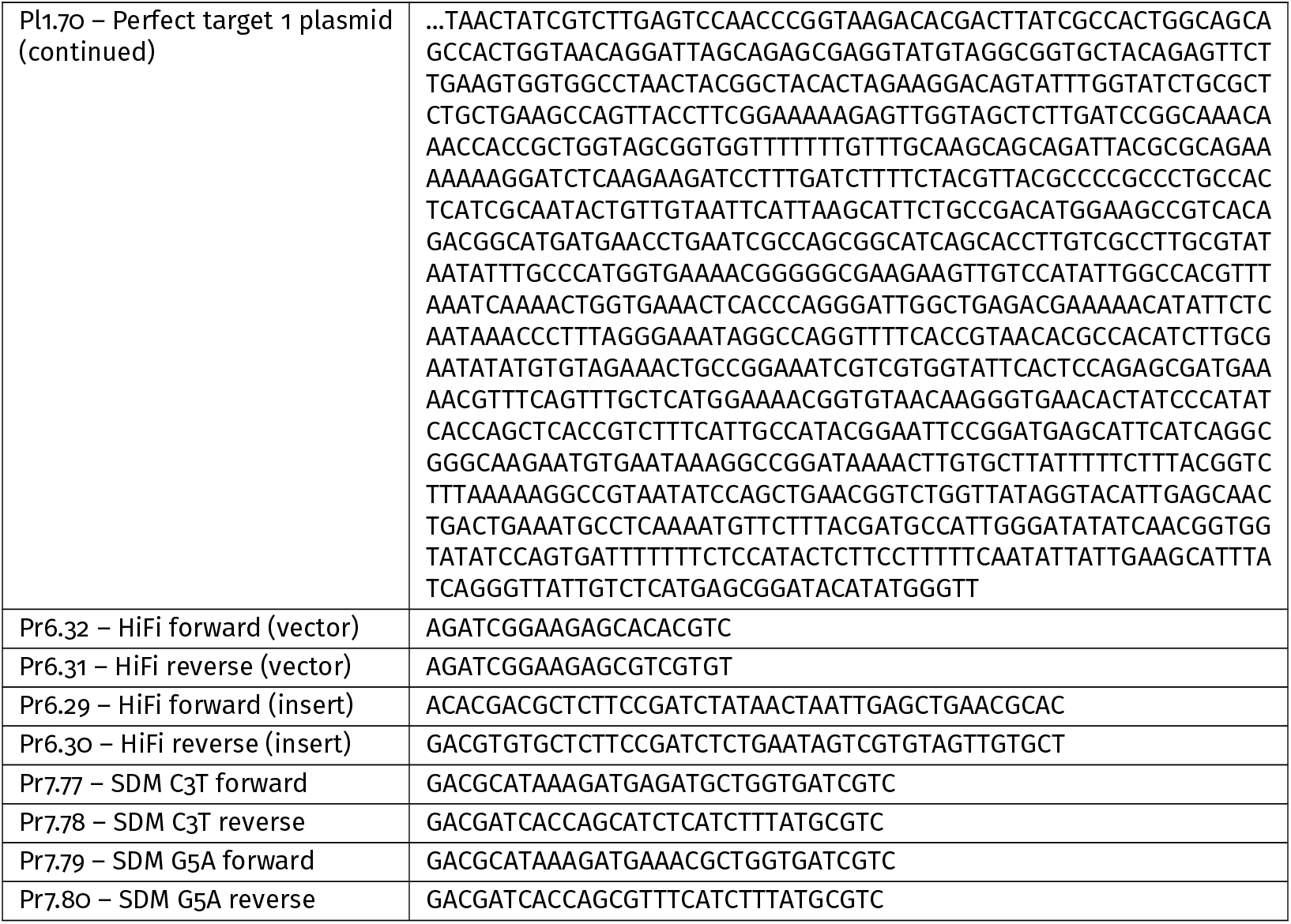
Plasmids, DNA targets and their primers used in the study.

## References

1. Wang, J. Y. & Doudna, J. A. CRISPR technology: A decade of genome editing is only the beginning. Science 379. Publisher: American Association for the Advancement of Science, eadd8643. doi:10.1126/science.add8643 (Jan. 2023).

2. Zhu, H., Li, C. & Gao, C. Applications of CRISPR–Cas in agriculture and plant biotechnology. en. Nature Reviews Molecular Cell Biology 21. Publisher: Nature Publishing Group, 661–677. issn: 1471-0080. doi:10.1038/s41580-020-00288-9 (Nov. 2020).

3. Morshedzadeh, F., Ghanei, M., Lotfi, M., Ghasemi, M., Ahmadi, M., Najari-Hanjani, P., Sharif, S., Mozaffari-Jovin, S., Peymani, M. & Abbaszadegan, M. R. An Update on the Application of CRISPR Technology in Clinical Practice. en. Molecular Biotechnology 66, 179–197. issn: 1559-0305. doi:10.1007/s12033-023-00724-z(Feb. 2024).

4. Chehelgerdi, M., Chehelgerdi, M., Khorramian-Ghahfarokhi, M., Shafieizadeh, M., Mahmoudi, E., Eskandari, F., Rashidi, M., Arshi, A. & Mokhtari-Farsani, A. Comprehensive review of CRISPR-based gene editing: mechanisms, challenges, and applications in cancer therapy. Molecular Cancer 23, 9. issn: 1476-4598. doi:10.1186/s12943-023-01925-5 (Jan. 2024).

5. Zahedipour, F., Zahedipour, F., Zamani, P., Jaafari, M. R. & Sahebkar, A. Harnessing CRISPR technology for viral therapeutics and vaccines: from preclinical studies to clinical applications. Virus Research 341, 199314. issn: 0168-1702. doi:10.1016/j.virusres.2024.199314 (Mar. 2024).

6. Jinek, M., Chylinski, K., Fonfara, I., Hauer, M., Doudna, J. A. & Charpentier, E. A programmable dual-RNA-guided DNA endonuclease in adaptive bacterial immunity. Science 337. Publisher: American Association for the Advancement of Science, 816–821. issn: 10959203. doi:10.1126/science.1225829 (Aug. 2012).

7. Scully, R., Panday, A., Elango, R. & Willis, N. A. DNA double-strand break repair-pathway choice in somatic mammalian cells. en. Nature Reviews Molecular Cell Biology 20. Publisher: Nature Publishing Group, 698–714. issn: 1471-0080. doi:10.1038/s41580-019-0152-0 (Nov. 2019).

8. Xue, C. & Greene, E. C. DNA Repair Pathway Choices in CRISPR-Cas9-Mediated Genome Editing. English. Trends in Genetics 37. Publisher: Elseviery639–656. issn: 0168-9525. doi:10.1016/j.tig.2021.02.008 (July 2021).

9. Zou, R. S., Marin-Gonzalez, A., Liu, Y., Liu, H. B., Shen, L., Dveirin, R. K., Luo, J. X. J., Kalhor, R. & Ha, T. Massively parallel genomic perturbations with multi-target CRISPR interrogates Cas9 activity and DNA repair at endogenous sites. en. Nature Cell Biology. issn: 1465-7392, 1476-4679. doi:10.1038/s41556-022-00975-z (Sept. 2022).

10. Moreb, E. A. & Lynch, M. D. Genome dependent Cas9/gRNA search time underlies sequence dependent gRNA activity. en. Nature Communications 12. Number: 1 Publisher: Nature Publishing Group, 5034. issn: 2041-1723. doi:10.1038/s41467-021-25339-3 (Aug. 2021).

11. Zhang, H., Yan, J., Lu, Z., Zhou, Y., Zhang, Q., Cui, T., Li, Y., Chen, H. & Ma, L. Deep sampling of gRNA in the human genome and deep-learning-informed prediction of gRNA activities. en. Cell Discovery 9. Publisher: Nature Publishing Group, 48. issn: 2056-5968. doi:10.1038/s41421-023-00549-9 (May 2023).

12. Cameron, P., Fuller, C. K., Donohoue, P. D., Jones, B. N., Thompson, M. S., Carter, M. M., Gradia, S., Vidal, B., Garner, E., Slorach, E. M., Lau, E., Banh, L. M., Lied, A. M., Edwards, L. S., Settle, A. H., Capurso, D., Llaca, V., Deschamps, S., Cigan, M., Young, J. K. & May, A. P. Mapping the genomic landscape of CRISPR–Cas9 cleavage. en. Nature Methods 14. Publisher: Nature Publishing Group, 600–606. issn: 1548-7105. doi:10.1038/nmeth.4284 (June 2017).

13. Tsai, S. Q., Nguyen, N. T., Malagon-Lopez, J., Topkar, V. V., Aryee, M. J. & Joung, J. K. CIRCLE-seq: a highly sensitive in vitro screen for genome-wide CRISPR–Cas9 nuclease off-targets. en. Nature Methods 14. Publisher: Nature Publishing Group, 607–614. issn: 1548-7105. doi:10.1038/nmeth.4278 (June 2017).

14. Fu, R., He, W., Dou, J., Villarreal, O. D., Bedford, E., Wang, H., Hou, C., Zhang, L., Wang, Y., Ma, D., Chen, Y., Gao, X., Depken, M. & Xu, H. Systematic decomposition of sequence determinants governing CRISPR/Cas9 specificity. Nature Communications 13. doi:10.1038/s41467-022-28028-x (2022).

15. Konstantakos, V., Nentidis, A., Krithara, A. & Paliouras, G. CRISPR–Cas9 gRNA efficiency prediction: an overview of predictive tools and the role of deep learning. Nucleic Acids Research 50, 3616–3637. issn: 0305-1048. doi:10.1093/nar/gkac192 (Apr. 2022).

16. Sharma, S., Murmu, S., Das, R., Tilgam, J., Saakre, M. & Paul, K. A review on bioinformatics advances in CRISPR-Cas technology. en. Journal of Plant Biochemistry and Biotechnology. issn: 0974-1275. doi:10.1007/s13562-022-00811-3 (Nov. 2022).

17. Chen, Y. & Wang, X. Evaluation of efficiency prediction algorithms and development of ensemble model for CRISPR/Cas9 gRNA selection. Bioinformatics 38, 5175– 5181. issn: 1367-4811. doi:10.1093/bioinformatics/btac681 (Dec. 2022).

18. Zhang, G., Luo, Y., Dai, X. & Dai, Z. Benchmarking deep learning methods for predicting CRISPR/Cas9 sgRNA on- and off-target activities. Briefings in Bioinformatics 24, bbad333. issn: 1477-4054. doi:10.1093/bib/bbad333 (Nov. 2023).

19. Yuan, H., Song, C., Xu, H., Sun, Y., Anthon, C., Bolund, L., Lin, L., Benabdellah, K., Lee, C., Hou, Y., Gorodkin, J. & Luo, Y. An Overview and Comparative Analysis of CRISPR-SpCas9 gRNA Activity Prediction Tools. The CRISPR Journal 8. Publisher: Mary Ann Liebert, Inc., publishers, 89–104. issn: 2573-1599. doi:10.1089/crispr.2024.0058 (Apr. 2025).

20. Haeussler, M., Schönig, K., Eckert, H., Eschstruth, A., Mianné, J., Renaud, J.-B., Schneider-Maunoury, S., Shkumatava, A., Teboul, L., Kent, J.Joly, J.-S. & Concordet, J.-P. Evaluation of off-target and on-target scoring algorithms and integration into the guide RNA selection tool CRISPOR. Genome Biology 17, 148. issn: 1474-760X. doi:10.1186/s13059-016-1012-2 (July 2016).

21. Eslami-Mossallam, B., Klein, M., Smagt, C. V. D., Sanden, K. V. D., Jones Jr., S.K., Hawkins, J. A., Finkelstein, I. J. & Depken, M. A kinetic model predicts SpCas9 activity, improves off-target classification, and reveals the physical basis of targeting fidelity. Nature Communications 13. doi:10.1038/s41467-022-28994-2 (2022).

22. Farasat, I. & Salis, H. M. A Biophysical Model of CRISPR/Cas9 Activity for Rational Design of Genome Editing and Gene Regulation. en. PLOS Computational Biology 12. Publisher: Public Library of Science, e1004724. issn: 1553-7358. doi:10.1371/journal.pcbi.1004724 (Jan. 2016).

23. Alkan, F., Wenzel, A., Anthon, C., Havgaard, J. H. & Gorodkin, J. CRISPR-Cas9 off-targeting assessment with nucleic acid duplex energy parameters. Genome Biology 19. Publisher: Genome Biology, 1–13. issn: 1474760X. doi:10.1186/s13059-018-1534-x (2018).

24. Zhang, D., Hurst, T., Duan, D. & Chen, S. J. Unified energetics analysis unravels SpCas9 cleavage activity for optimal gRNA design. Proceedings of the National Academy of Sciences of the United States of America 116. Publisher: National Academy of Sciences, 8693– 8698. issn: 10916490. doi:10.1073/pnas.1820523116 (Apr. 2019).

25. Corsi, G. I., Qu, K., Alkan, F., Pan, X., Luo, Y. & Gorodkin, J. CRISPR/Cas9 gRNA activity depends on free energy changes and on the target PAM context. en. Nature Communications 13. Number: 1 Publisher: Nature Publishing Group, 3006. issn: 2041-1723. doi:10.1038/s41467-022-30515-0 (May 2022).

26. Bisaria, N., Jarmoskaite, I. & Herschlag, D. Lessons from Enzyme Kinetics Reveal Specificity Principles for RNA-Guided Nucleases in RNA Interference and CRISPR-Based Genome Editing. Cell Systems 4, 21–29. issn: 2405-4712. doi:10.1016/j.cels.2016.12.010 (Jan. 2017).

27. Klein, M., Eslami-Mossallam, B., Arroyo, D. G. & Depken, M. Hybridization Kinetics Explains CRISPR-Cas Off-Targeting Rules. Cell Reports 22. Publisher: Elsevier, 1413–1423. issn: 22111247. doi:10.1016/j.celrep.2018.01.045 (Feb. 2018).

28. Ratajczyk, E. J., Bath, J., Šulc, P., Doye, J. P. K., Louis, A. & Turberfield, A. J. Controlling DNA–RNA strand displacement kinetics with base distribution. Proceedings of the National Academy of Sciences 122. Publisher: Proceedings of the National Academy of Sciences, e2416988122. doi:10.1073/pnas.2416988122 (June 2025).

29. Anders, C., Niewoehner, O., Duerst, A. & Jinek, M. Structural basis of PAM-dependent target DNA recognition by the Cas9 endonuclease. en. Nature 513. Publisher: Nature Publishing Group, 569–573. issn: 1476-4687. doi:10.1038/nature13579 (Sept. 2014).

30. Nishimasu, H., Ran, F. A., Hsu, P. D., Konermann, S., Shehata, S.I., Dohmae, N., Ishitani, R., Zhang, F. & Nureki, O. Crystal Structure of Cas9 in Complex with Guide RNA and Target DNA. Cell 156, 935–949. issn: 0092-8674. doi:10.1016/j.cell.2014.02.001 (Feb. 2014).

31. Jiang, F., Zhou, K., Ma, L., Gressel, S. & Doudna, J. A. A Cas9-guide RNA complex preorganized for target DNA recognition. Science 348. Publisher: American Association for the Advancement of Science, 1477–1481. issn: 10959203. doi:10.1126/science.aab1452 (June 2015).

32. Pacesa, M., Loeff, L., Querques, I., Muckenfuss, L. M., Sawicka, M. & Jinek, M. R-loop formation and conformational activation mechanisms of Cas9. en. Nature 609. Number: 7925 Publisher: Nature Publishing Group, 191–196. issn: 1476-4687. doi:10.1038/s41586-022-05114-0 (Sept. 2022).

33. Rutkauskas, M., Sinkunas, T., Songailiene, I., Tikhomirova, M. S., Siksnys, V. & Seidel, R. Directional R-loop formation by the CRISPR-cas surveillance complex cascade provides efficient off-target site rejection. Cell Reports 10. Publisher: Elsevier, 1534–1543. issn: 22111247. doi:10.1016/j.celrep.2015.01.067 (Mar. 2015).

34. Singh, D., Sternberg, S. H., Fei, J., Doudna, J. A. & Ha, T. Real-time observation of DNA recognition and rejection by the RNA-guided endonuclease Cas9. Nature Communications 7. Publisher: Nature Publishing Group, 1–8. issn: 20411723. doi:10.1038/ncomms12778 (Sept. 2016).

35. Jiang, F., Taylor, D. W., Chen, J. S., Kornfeld, J. E., Zhou, K., Thompson, A. J., Nogales, E. & Doudna, J. A. Structures of a CRISPR-Cas9 R-loop complex primed for DNA cleavage. Science 351. Publisher: American Association for the Advancement of Science, 867–871. issn: 10959203. doi:10.1126/science.aad8282 (Feb. 2016).

36. Zhu, X., Clarke, R., Puppala, A. K., Chittori, S., Merk, A., Merrill, B. J., Simonović, M. & Subramaniam, S. Cryo-EM structures reveal coordinated domain motions that govern DNA cleavage by Cas9. en. Nature Structural & Molecular Biology 26. Publisher: Nature Publishing Group, 679–685. issn: 1545-9985. doi:10.1038/s41594-019-0258-2 (Aug. 2019).

37. Gong, S., Yu, H. H., Johnson, K. A. & Taylor, D. W. DNA Unwinding Is the Primary Determinant of CRISPR-Cas9 Activity. Cell Reports 22. Publisher: Elsevier B.V., 359– 371. issn: 22111247. doi:10.1016/J.CELREP.2017.12.041 (2018).

38. Raper, A. T., Stephenson, A. A. & Suo, Z. Functional Insights Revealed by the Kinetic Mechanism of CRISPR/Cas9. Journal of the American Chemical Society 140. Publisher: American Chemical Society, 2971– 2984. issn: 15205126. doi:10.1021/jacs.7b13047 (Feb. 2018).

39. SantaLucia, J. & Hicks, D. The Thermodynamics of DNA Structural Motifs. en. Annual Review of Biophysics 33. Publisher: Annual Reviews, 415–440. issn: 1936-122X, 1936-1238. doi:10.1146/annurev.biophys.32.110601.141800 (June 2004).

40. Sugimoto, N., Nakano, S.-i., Katoh, M., Matsumura, A., Nakamuta, H., Ohmichi, T., Yoneyama, M. & Sasaki, M. Thermodynamic Parameters To Predict Stability of RNA/DNA Hybrid Duplexes. Biochemistry 34. Publisher: American Chemical Society, 11211–11216. issn: 0006-2960. doi:10.1021/bi00035a029 (Sept. 1995).

41. Sugimoto, N., Nakano, M. & Nakano, S.-i. Thermodynamics-Structure Relationship of Single Mismatches in RNA/DNA Duplexes. Biochemistry 39. Publisher: American Chemical Society, 11270–11281. issn: 0006-2960. doi:10.1021/bi000819p (Sept. 2000).

42. Watkins Jr, N. E., Kennelly, W. J., Tsay, M. J., Tuin, A., Swenson, L., Lee, H.-R., Morosyuk, S., Hicks, D. A. & SantaLucia Jr, J. Thermodynamic contributions of single internal rA·dA, rC·dC, rG·dG and rU·dT mismatches in RNA/DNA duplexes. Nucleic Acids Research 39, 1894– 1902. issn: 0305-1048. doi:10.1093/nar/gkq905 (Mar. 2011).

43. Turner, D. H. & Mathews, D. H. NNDB: the nearest neighbor parameter database for predicting stability of nucleic acid secondary structure. Nucleic Acids Research 38, D280–D282. issn: 0305-1048. doi:10.1093/nar/gkp892 (Jan. 2010).

44. Nakano, S.-i., Fujimoto, M., Hara, H. & Sugimoto, N. Nucleic acid duplex stability: influence of base composition on cation effects. Nucleic Acids Research 27, 2957– 2965. issn: 0305-1048. doi:10.1093/nar/27.14.2957 (July 1999).

45. Banerjee, D., Tateishi-Karimata, H., Ohyama, T., Ghosh, S., Endoh, T., Takahashi, S. & Sugimoto, N. Improved nearest-neighbor parameters for the stability of RNA/DNA hybrids under a physiological condition. Nucleic Acids Research 48, 12042–12054. issn: 0305-1048. doi:10.1093/nar/gkaa572 (Dec. 2020).

46. Sternberg, S. H., Redding, S., Jinek, M., Greene, E. C. & Doudna, J. A. DNA interrogation by the CRISPR RNA-guided endonuclease Cas9. Nature 507. Publisher: Nature Publishing Group, 62–67. issn: 14764687. doi:10.1038/nature13011 (Jan. 2014).

47. Ivanov, I. E., Wright, A. V., Cofsky, J. C., Palacio Aris, K. D., Doudna, J. A. & Bryant, Z. Cas9 interrogates DNA in discrete steps modulated by mismatches and supercoiling. Proceedings of the National Academy of Sciences of the United States of America 117, 5853–5860. issn: 10916490. doi:10.1073/pnas.1913445117 (2020).

48. Boyle, E. A., Andreasson, J. O. L., Chircus, L. M., Sternberg, S.H., Wu, M. J., Guegler, C. K., Doudna, J. A. & Greenleaf, W. J. High-throughput biochemical profiling reveals sequence determinants of dCas9 off-target binding and unbinding. en. Proceedings of the National Academy of Sciences 114, 5461–5466. issn: 0027-8424, 1091-6490. doi:10.1073/pnas.1700557114 (May 2017).

49. Aguirre Rivera, J., Mao, G., Sabantsev, A., Panfilov, M., Hou, Q., Lindell, M., Chanez, C., Ritort, F., Jinek, M. & Deindl, S. Massively parallel analysis of single-molecule dynamics on next-generation sequencing chips. Science 385. Publisher: American Association for the Advancement of Science, 892–898. doi:10.1126/science.adn5371 (Aug. 2024).

50. Newton, M. D., Losito, M., Smith, Q. M., Parnandi, N., Taylor, B. J., Akcakaya, P., Maresca, M., van Eijk, P., Reed, S. H., Boulton, S. J., King, G. A., Cuomo, M. E. & Rueda, D. S. Negative DNA supercoiling induces genome-wide Cas9 off-target activity. Molecular Cell 83, 3533–3545.e5. issn: 1097-2765. doi:10.1016/j.molcel.2023.09.008 (Oct. 2023).

51. Banerjee, T., Takahashi, H., Rendra, D., Subekti, G. & Kamagata, K. Engineering of the genome editing protein Cas9 to slide along DNA. Scientific Reports 11. ISBN: 0123456789, 14165. doi:10.1038/s41598-021-93685-9(2021).

52. Ratajczyk, E. J., Šulc, P., Turberfield, A. J., Doye, J. P. K. & Louis, A. A. Coarse-grained modeling of DNA–RNA hybrids. The Journal of Chemical Physics 160, 115101. issn: 0021-9606. doi:10.1063/5.0199558 (Mar. 2024).

53. Chen, Q., Chuai, G., Zhang, H., Tang, J., Duan, L., Guan, H., Li, W., Li, W., Wen, J., Zuo, E., Zhang, Q. & Liu, Q. Genome-wide CRISPR off-target prediction and optimization using RNA-DNA interaction fingerprints. en. Nature Communications 14. Publisher: Nature Publishing Group, 7521. issn: 2041-1723. doi:10.1038/s41467-023-42695-4 (Nov. 2023).

54. Cofsky, J. C., Soczek, K. M., Knott, G. J., Nogales, E. & Doudna, J. A. CRISPR–Cas9 bends and twists DNA to read its sequence. Nature Structural & Molecular Biology 29, 395–402. issn: 1545-9993. doi:10.1038/s41594-022-00756-0 (Apr. 2022).

55. Bravo, J. P. K., Liu, M.-S., Hibshman, G. N., Dangerfield, T. L., Jung, K., McCool, R. S., Johnson, K. A. & Taylor, D. W. Structural basis for mismatch surveillance by CRISPR–Cas9. Nature. issn: 0028-0836. doi:10.1038/S41586-022-04470-1 (Mar. 2022).

56. Boyle, E. A., Becker, W. R., Bai, H. B., Chen, J. S., Doudna, J. A. & Greenleaf, W. J. Quantification of Cas9 binding and cleavage across diverse guide sequences maps landscapes of target engagement. Science Advances 7. issn: 23752548. doi:10.1126/sciadv.abe5496 (2021).

57. Rutkauskas, M., Songailiene, I., Irmisch, P., Kemmerich, F. E., Sinkunas, T., Siksnys, V. & Seidel, R. A quantitative model for the dynamics of target recognition and off-target rejection by the CRISPR-Cas Cascade complex. en. Nature Communications 13. Number: 1 Publisher: Nature Publishing Group, 7460. issn: 2041-1723. doi:10.1038/s41467-022-35116-5 (Dec. 2022).

58. Aris, K. D. P., Cofsky, J. C., Shi, H., Al-Sayyad, N., Ivanov, I. E., Balaji, A., Doudna, J. A. & Bryant, Z. Dynamic basis of supercoiling-dependent DNA interrogation by Cas12a via R-loop intermediates. en. Nature Communications 16. Publisher: Nature Publishing Group, 2939. issn: 2041-1723. doi:10.1038/s41467-025-57703-y (Mar. 2025).

59. Shi, H., Al-Sayyad, N., Wasko, K. M., Trinidad, M. I., Doherty, E. E., Vohra, K., Boger, R. S., Colognori, D., Cofsky, J. C., Skopintsev, P., Bryant, Z. & Doudna, J. A. Rapid two-step target capture ensures efficient CRISPR-Cas9-guided genome editing. Molecular Cell 85, 1730–1742.e9. issn: 1097-2765. doi:10.1016/j.molcel.2025.03.024 (May 2025).

60. Kauert, D. J., Madariaga-Marcos, J., Rutkauskas, M., Wulfken, A., Songailiene, I., Sinkunas, T., Siksnys, V. & Seidel, R. The energy landscape for R-loop formation by the CRISPR–Cas Cascade complex. en. Nature Structural & Molecular Biology 30. Number: 7 Publisher: Nature Publishing Group, 1040–1047. issn: 1545-9985. doi:10.1038/s41594-023-01019-2 (July 2023).

61. Pacesa, M., Lin, C.-H., Cléry, A., Saha, A., Arantes, P. R., Bargsten, K., Irby, M. J., Allain, F. H. .-., Palermo, G., Cameron, P., Donohoue, P. D. & Jinek, M. Structural basis for Cas9 off-target activity. Cell 185, 4067–4081.e21. issn: 0092-8674. doi:10.1016/j.cell.2022.09.026 (Oct. 2022).

62. Leenay, R. T., Maksimchuk, K. R., Slotkowski, R. A., Agrawal, R. N., Gomaa, A. A., Briner, A. E., Barrangou, R. & Beisel, C. L. Identifying and Visualizing Functional PAM Diversity across CRISPR-Cas Systems. Molecular Cell 62. Publisher: Elsevier Inc., 137–147. issn: 10974164. doi:10.1016/j.molcel.2016.02.031 (2016).

63. Jones Jr., S.K., Hawkins, J. A., Johnson, N. V., Jung, C., Hu, K., Rybarski, J. R., Chen, J. S., Doudna, J. A., Press, W. H. & Finkelstein, I. J. Massively parallel kinetic profiling of natural and engineered CRISPR nucleases. en. Nature Biotechnology 39. Number: 1 Publisher: Nature Publishing Group, 84–93. issn: 1546-1696. doi:10.1038/s41587-020-0646-5 (Jan. 2021).

64. Lu, Q., Bhat, D., Stepanenko, D. & Pigolotti, S. Search and Localization Dynamics of the CRISPR-Cas9 System. Physical Review Letters 127. Publisher: American Physical Society, 208102. doi:10.1103/PhysRevLett.127.208102 (Nov. 2021).

65. Achar, Y. J., Adhil, M., Choudhary, R., Gilbert, N. & Foiani, M. Negative supercoil at gene boundaries modulates gene topology. en. Nature 577. Publisher: Nature Publishing Group, 701–705. issn: 1476-4687. doi:10.1038/s41586-020-1934-4 (Jan. 2020).

66. Jaskovikaitė, I., Offerhaus, H. S., Vinogradovas, M., Barkauskaitė, U., Depken, M. & Jones Jr., S.K. Supercoiling twists Cas9 off-target discrimination when binding, nicking and cleaving. Manuscript in preparation.

67. Raguram, A., Banskota, S. & Liu, D. R. Therapeutic in vivo delivery of gene editing agents. English. Cell 185. Publisher: Elsevier, 2806–2827. issn: 0092-8674, 1097-4172. doi:10.1016/j.cell.2022.03.045 (July 2022).

68. Karp, H., Zoltek, M., Wasko, K., Vazquez, A. L., Brim, J., Ngo, W., Schepartz, A. & Doudna, J. Packaged delivery of CRISPR-Cas9 ribonucleoproteins accelerates genome editing. eng. bioRxiv: The Preprint Server for Biology, 2024.10.18.619117. issn: 2692-8205. doi:10.1101/2024.10.18.619117 (Oct. 2024).

69. Moreb, E. A., Hutmacher, M. & Lynch, M. D. CRISPR-Cas “Non-Target” Sites Inhibit On-Target Cutting Rates. The CRISPR Journal 3. Publisher: Mary Ann Liebert, Inc., publishers, 550–561. issn: 2573-1599. doi:10.1089/crispr.2020.0065 (Dec. 2020).

70. Offerhaus, H. CRISPRzip-tool version v1.0.0. Dec. 2025. doi:10.5281/zenodo.17818523.

71. Offerhaus, H. CRISPRzip version 1.2.2. Dec. 2025. doi:10.5281/zenodo.17818249.

72. Zhang, Z., Lamson, A. R., Shelley, M. & Troyanskaya, O. Interpretable neural architecture search and transfer learning for understanding CRISPR–Cas9 off-target enzymatic reactions. en. Nature Computational Science 3. Publisher: Nature Publishing Group, 1056–1066. issn: 2662-8457. doi:10.1038/s43588-023-00569-1 (Dec. 2023).

73. Knott, G. J. & Doudna, J. A. CRISPR-Cas guides the future of genetic engineering. Science 361, 866–869. issn: 10959203. doi:10.1126/science.aat5011 (2018).

74. Huang, X., Yang, D., Zhang, J., Xu, J. & Chen, Y. E. Recent Advances in Improving Gene-Editing Specificity through CRISPR–Cas9 Nuclease Engineering. en. Cells 11. Number: 14 Publisher: Multidisciplinary Digital Publishing Institute, 2186. issn: 2073-4409. doi:10.3390/cells11142186 (Jan. 2022).

75. Collias, D. & Beisel, C. L. CRISPR technologies and the search for the PAM-free nuclease. en. Nature Communications 12. Publisher: Nature Publishing Group, 555. issn: 2041-1723. doi:10.1038/s41467-020-20633-y (Jan. 2021).

76. Hawkins, J. A., Jones Jr., S.K., Finkelstein, I. J. & Press, W. H. Indel-correcting DNA barcodes for high-throughput sequencing. Proceedings of the National Academy of Sciences 115. Publisher: Proceedings of the National Academy of Sciences, E6217–E6226. doi:10.1073/pnas.1802640115 (July 2018).

77. Tsallis, C. & Stariolo, D. A. Generalized simulated annealing. en. Physica A: Statistical Mechanics and its Applications 233, 395–406. issn: 0378-4371. doi:10.1016/S0378-4371(96)00271-3 (Nov. 1996).

78. (DHPC), D. H. P. C. C. DelftBlue Supercomputer (Phase 2) https://www.tudelft.nl/dhpc/ark:/44463/DelftBluePhase2. https://www.tudelft.nl/dhpc/ark:/44463/DelftBluePhase2. 2024.

79. Marko, J. F. en. in Mathematics of DNA Structure, Function and Interactions 1st ed. The IMA Volumes in Mathematics and its Applications 150, 225–249 (Springer, New York, NY, 2020). isbn: 978-1-4419-0669-4.

80. Gao, X., Hong, Y., Ye, F., Inman, J. T. & Wang, M. D. Torsional Stiffness of Extended and Plectonemic DNA. en. Physical Review Letters 127. Publisher: American Physical Society, 028101. doi:10.1103/PhysRevLett.127.028101 (July 2021).

